# Population genomics of bank vole populations reveals associations between immune related genes and the epidemiology of Puumala hantavirus in Sweden

**DOI:** 10.1101/148163

**Authors:** Audrey Rohfritsch, Maxime Galan, Mathieu Gautier, Karim Gharbi, Gert Olsson, Bernhard Gschloessl, Caroline Zeimes, Sophie VanWambeke, Renaud Vitalis, Nathalie Charbonnel

## Abstract

Infectious pathogens are major selective forces acting on individuals. The recent advent of high-throughput sequencing technologies now enables to investigate the genetic bases of resistance/susceptibility to infections in non-model organisms. From an evolutionary perspective, the analysis of the genetic diversity observed at these genes in natural populations provides insight into the mechanisms maintaining polymorphism and their epidemiological consequences. We explored these questions in the context of the interactions between Puumala hantavirus (PUUV) and its reservoir host, the bank vole *Myodes glareolus*. Despite the continuous spatial distribution of *M. glareolus* in Europe, PUUV distribution is strongly heterogeneous. Different defence strategies might have evolved in bank voles as a result of co-adaptation with PUUV, which may in turn reinforce spatial heterogeneity in PUUV distribution. We performed a genome scan study of six bank vole populations sampled along a North/South transect in Sweden, including PUUV endemic and non-endemic areas. We combined candidate gene analyses (*Tlr4*, *Tlr7*, *Mx2* genes) and high throughput sequencing of RAD (Restriction-site Associated DNA) markers. We found evidence for outlier loci showing high levels of genetic differentiation. Ten outliers among the 52 that matched to mouse protein-coding genes corresponded to immune related genes and were detected using ecological associations with variations in PUUV prevalence. One third of the enriched pathways concerned immune processes, including platelet activation and TLR pathway. In the future, functional experimentations should enable to confirm the role of these these immune related genes with regard to the interactions between *M. glareolus* and PUUV.

## Introduction

Infections are among the strongest selective forces acting in natural populations (Fumagalli et al., 2011; Karlsson, Kwiatkowski, & Sabeti, 2014). As a consequence, hosts have evolved a wide range of defence mechanisms against their pathogens. Understanding variation in host defence mechanisms has been at the core of eco-immunology in the last two decades (Sheldon & Verhulst, 1996). This immunoheterogeneity seems to be strongly driven by non-heritable influences (e.g., age, physiological status, resource availability, microbial exposure, history of infections during lifetime, Schmid-Hempel, 2003; Schulenburg, Kurtz, Moret, & Siva-Jothy, 2009). It also has a genetic basis that is highly correlated to immune-related genes (Barreiro & Quintana-Murci, 2010; Hill et al., 1994). In this context, population genetics approaches may help deciphering the relative influence of different evolutionary processes (including migration, drift, and selection) in shaping the variation of immune-related genes in space and time (Charbonnel & Cosson, 2011; Quintana-Murci & Clark, 2013). In turn, the analysis of the genetic diversity observed at immune-related genes in natural populations enables to elucidate some geographical patterns of pathogen distribution. Still, only few studies address the question of how pathogen-mediated selective processes in the hosts may shape the spatial distribution of pathogens but see (Guivier, Galan, Henttonen, Cosson, & Charbonnel, 2014; Guivier et al., 2010; Wenzel, Douglas, James, Redpath, & Piertney, 2016).

A particularly relevant study system to tackle these questions consists of the interaction between the hantavirus Puumala (PUUV), the causative agent of nephropathia epidemica in humans (Brummer-Korvenkotio, Henttonen, & Vaheri, 1982), and its reservoir host, the bank vole *Myodes glareolus*. Despite the continuous spatial distribution of *M. glareolus* in Europe (Stenseth, 1985), PUUV distribution is strongly heterogeneous (Olsson, Leirs, & Henttonen, 2010). Different hypotheses have been sought to explain this discrepancy. First, environmental variables that affect bank vole population size, e.g. landscape fragmentation or low snow cover, are likely to affect PUUV epidemiology and prevent PUUV persistence within reservoir host populations. They may at least partly explain the absence of PUUV in particular geographic areas. Second, environmental factors may also affect the possibility for PUUV to survive outside its reservoir host (e.g. low soil humidity or high temperatures, Sauvage, Langlais, Yoccoz, & Pontier, 2003). However, ecological niche modelling based on these environmental variables failed to explain accurately the distribution of PUUV in Europe (Zeimes, Olsson, Ahlm, & Vanwambeke, 2012; Zeimes et al., 2015). An alternative hypothesis states that spatial variation in the outcomes of *M. glareolus* / PUUV interactions may affect PUUV replication and excretion in the environment, which could ultimately shape PUUV distribution and nephropathia epidemica incidence in Europe (Guivier et al., 2014; Rohfritsch, Guivier, Galan, Chaval, & Charbonnel, 2013). Indeed, voles differ in their probability of being infected by PUUV (Kallio et al., 2006) and experimental infections have confirmed that the outcome of PUUV infection could vary between individuals (Dubois, Castel, et al., 2017; Hardestam et al., 2008). Furthermore, the genetic background of bank voles contributes to the variation in the response to infection (Charbonnel et al., 2014). In particular, differences in SNP allele frequencies within the tumor necrosis factor (*Tnf*) promoter and the *Mx2* gene are likely to influence PUUV distribution and epidemiology (Guivier et al., 2014; Guivier et al., 2010).

So far, the role of immune-related genes on bank vole response to PUUV infection has only been investigated for a handful of candidate genes (Charbonnel et al., 2014). The recent advent of high-throughput sequencing technologies now offers the opportunity to test for associations between genetic polymorphisms and susceptibility to infections in natural populations, at a genome-wide scale. Here, we focus on bank vole populations from Sweden, a relevant geographic area for the purpose of this study since the distribution of PUUV is highly heterogeneous throughout the country. Nephropathia epidemica is endemic in the central and northern parts of the country (Niklasson & LeDuc, 1987), with about 90% of all human cases in Sweden being found in the four northernmost counties. In particular, Västerbotten county exhibits the highest nephropathia epidemica incidence in Sweden, and probably even worldwide (Petterson, Boman, Juto, Evander, & Ahlm, 2008). This geographic pattern is not explained by the reservoir distribution because the bank vole is also common in the South of Sweden (Hörling et al., 1996). Furthermore, Sweden is characterized by a wide range of climatic and ecological conditions, which influence the spatial vegetation pattern. In particular, the landscape forest is divided in several vegetation zones, from the nemoral zone in the south where broad-leved deciduous forests dominate, the hemiboreal transition with mixed deciduous and coniferous forests to the northern wide belt of boreal forests, and up to the arctic tundra in the north. Finally, there is a contact zone at around 63° N in Sweden where two subpopulations of *M. glareolus* characterized by differentiated mitochondrial lineages meet (Nemirov, Leirs, Lundkvist, & Olsson, 2010). It is assumed that this contact zone was established during the Late Weichselian deglaciation, when bank voles re-colonized the Scandinavian Peninsula through a first, South-Scandinavian, migration stream across a pre-historic land bridge from present Denmark, and a second migration stream from the North-East (Tegelstrom, 1987). It has been shown that the PUUV strains present in the northern bank vole population are differentiated from those circulating in the southern bank vole populations (Hörling et al., 1996; Johansson et al., 2008).

The main objective of the present study was to characterize genome-wide patterns of bank vole population differentiation along a North/South transect in Sweden, and to identify specific genomic regions showing footprints of divergent selection between PUUV endemic areas in the North and non-endemic areas in the South. To that end, we used a population genomics approach relying on the sequencing of restriction-site-associated DNA (RAD-seq, see Baird et al., 2008) of pools of DNA from individuals sampled in six different localities and that have previously been characterized for a set of candidate genes (Dubois, Galan, et al., 2017). We combined different model-based methods of genome-scan, that allowed us to consider several underlying demographic scenarios, as well as putative associations with environmental variables. Last, we specifically asked whether immune-related genes were overrepresented among the genomic regions identified as presumably targeted by selection, as expected under the hypothesis that bank vole defence strategies against PUUV may have evolved differently in PUUV endemic and non-endemic areas. Overall, our study provides new insights into the selective processes that are likely to be involved in *M. glareolus* / PUUV interactions, and the mechanisms underlying the defence strategies of *M. glareolus* against PUUV infections. As such, it contributes to a better understanding of the factors driving PUUV distribution in reservoir populations, which is an important pre-requisite to apprehend the risk of nephropathia epidemica in Sweden.

## Materials and methods

### Sampling

Sampling was performed in April and October 2012. Using snap trapping, a total of 257 bank voles were caught in six localities distributed along a transect running from the North of Sweden, which is known to be highly endemic for PUUV, to the South of Sweden, where PUUV is absent in both bank voles and humans (Fig. 1, Table 1). Collected voles were kept on ice and transferred to -20°C freezers, before being processed in the laboratory. A piece of hind foot was placed in 95 % ethanol for further analyses. Permission to trap voles was obtained from the Swedish Environmental Protection Agency (SEPA; latest permission: Dnr 412-4009-10) and from the Animal Ethics Committee in Umeå (latest permission: Dnr A-61-11).

**Fig. 1.**
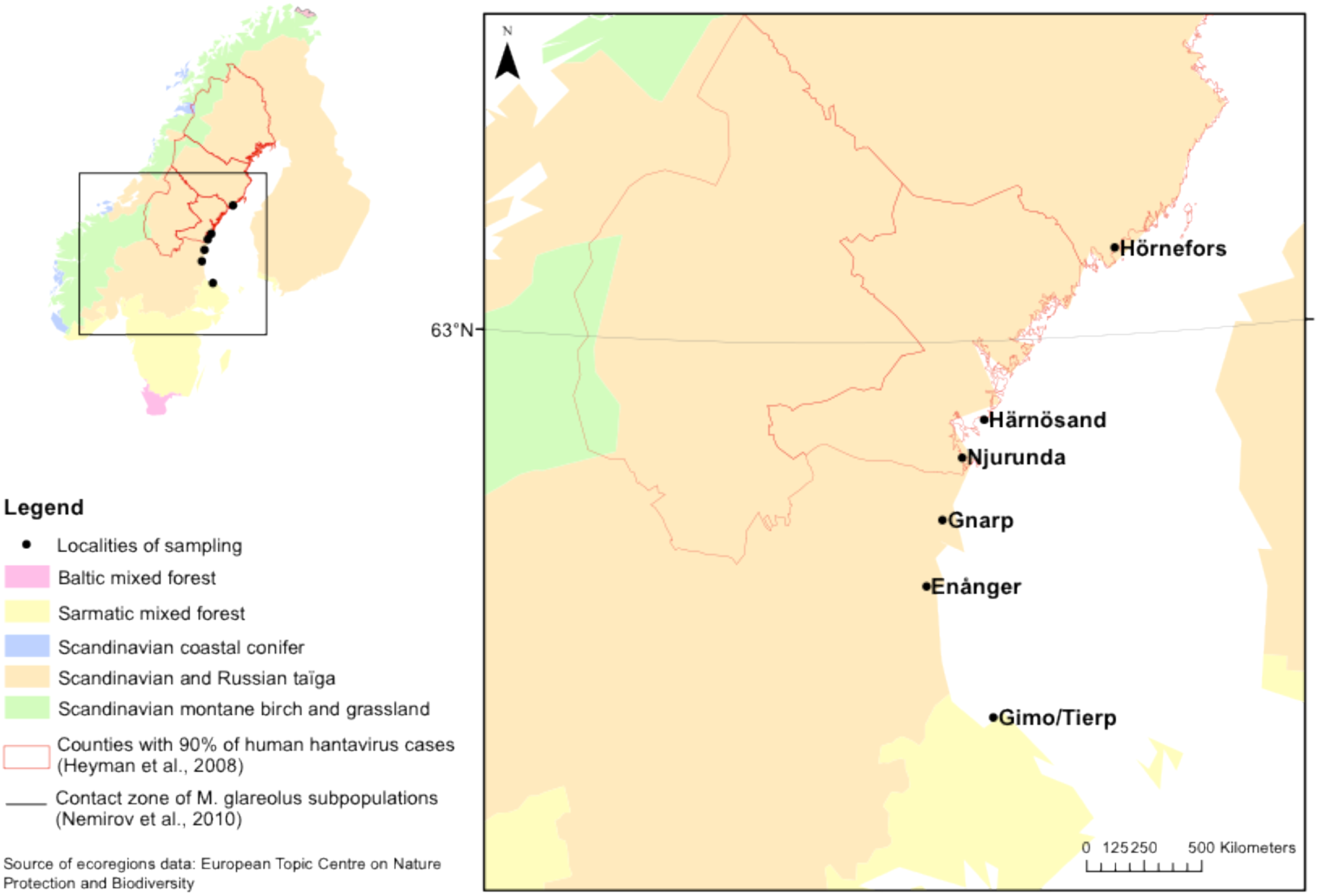
Maps showing the localities of bank vole sampling in Sweden. The red lines delimit the counties with 90% of human hantavirus cases reported. Geographic variations in ecoregions are represented with yellow, orange and green colors. The contact zone between the two mitochondrial lineages of *M. glareolus* is indicated with a black line.

**Table 1.**
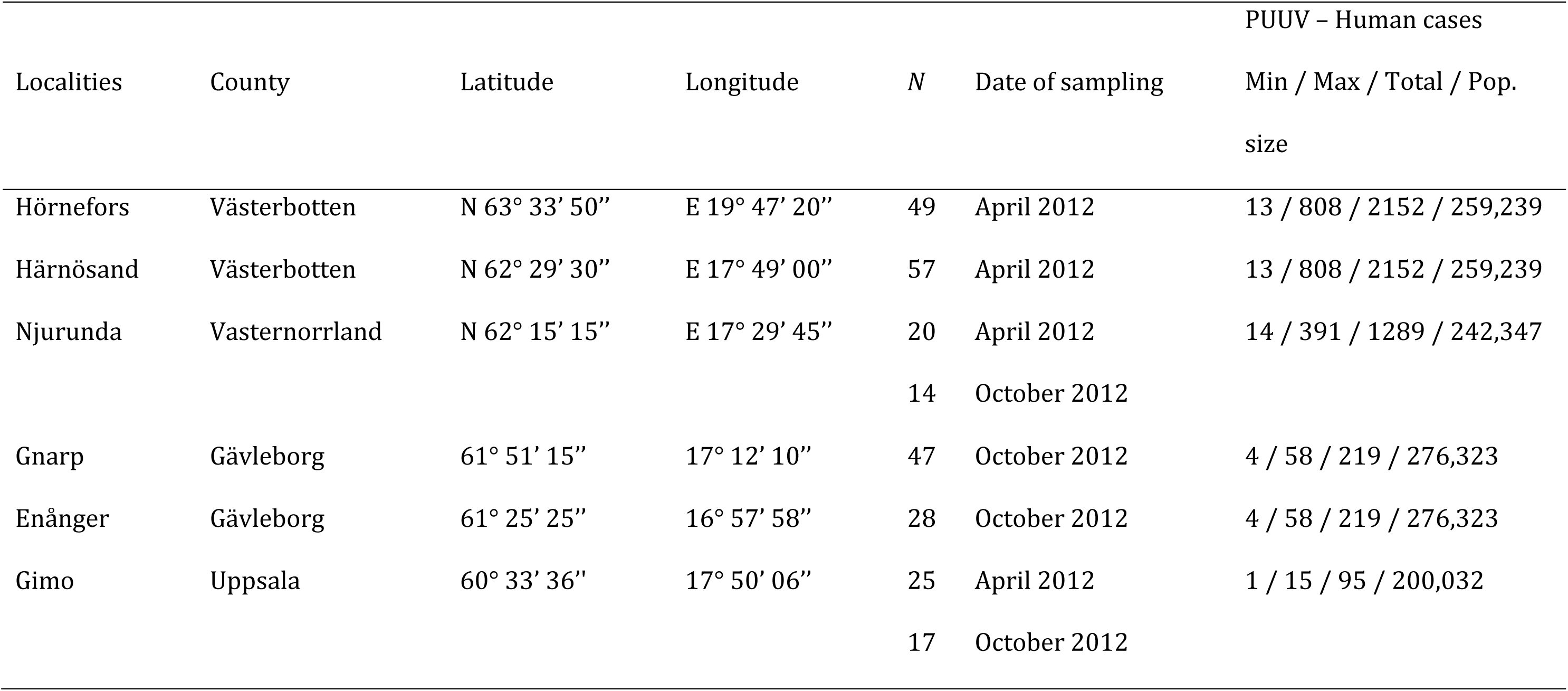
Sampling information, including localities of sampling and their administrative county, geographic coordinates (centre point from where the voles are trapped, voles are all caught within 1 kilometer from that point), the number of voles trapped *N*, the date of sampling, and the minimum, maximum and total number of human cases reported per county between 2001 and 2011 (SMI data).

### Molecular markers

Genomic DNA was extracted using the EZ-10 96-well plate genomic DNA isolation Kit for animal samples (Bio Basic Inc.) following the manufacturer recommendations.

#### Immune-related candidate genes

We studied the polymorphism of immune-related genes that have previously been shown to be associated with PUUV infections (Tnf promoter, Mx2, Tlr4 and Tlr7 genes, see for a review Charbonnel et al., 2014). The detection of polymorphisms in *Tnf* promoter was assessed using primers and PCR conditions derived from Guivier et al. (2010). For all other candidate genes, sequences of rat and mouse (*Mx2* exons 13 and 14 cDNA, *Tlr4* cDNA, and *Tlr7* cDNA) were retrieved from the Ensembl website (see Suppl. Mat. Table S1 for accession numbers). Specific primers were developed for all immune genes in Primer Designer - version 2.0 (Scientific and educational Software Program 1991). Because the exon 3 of *Tlr*s could not be sequenced at once (2251bp for *Tlr4*, 3149bp for *Tlr7*), we designed two primer sets to sequence two fragments of about 1000 bp for each *Tlr* (see Suppl. Mat. Table S2 for primer sequences). We first assessed polymorphism based on 12 bank voles. PCRs were performed on an Eppendorf Mastercycler EPgradient S (Hamburg, Germany) in a 25μl volume containing 0.5μl of each primer (10μM), 12.5μl of 2x Qiagen Multiplex PCR Master Mix, 9μl of ultrapure water and 2.5μl of extracted DNA. The cycling conditions included an initial denaturation at 95°C (15min) followed by a touchdown on the 10 first cycles of denaturation at 94°C (20s *Tnf*; 40s *Mx2* and *Tlrs*), annealing at 65–55°C (30s *Tnf*: 45s *Mx2* and *Tlrs*) with one degree less at each cycle, extension at 72°C (60s *Tnf*, 45s *Mx2*, 90s *Tlrs*), then 30 cycles of denaturation at 94°C (20s *Tnf*; 40s *Mx2* and *Tlrs*), annealing at 57°C (30s) for *Tnf* or 55°C (45s) for *Mx2* and *Tlrs* and extension at 72°C (60s *Tnf*, 45s *Mx2*, 90s *Tlrs*), and a final elongation step at 72°C for 10min.

The products of all these reactions were electrophoresed on 1.5% agarose gels and visualized by ethidium bromide staining. PCR products were sequenced in both directions by Eurofins MWG Operon (Ebersberg, Germany). Sequences were edited and aligned in BioEdit Sequences Alignment Editor using ClustalW Multiple Alignment (Hall 1999). We next used KASP genotyping services from LGC company (Genotyping by Allele-Specific Amplification Cuppen 2007) to genotype all sampled voles at SNPs that were polymorphic in the Swedish samples.

#### RAD tag sequencing

We chose to develop high throughput sequencing pools of individuals to reduce sequencing efforts and costs (Gautier et al., 2013; Schlötterer, Tobler, Kofler, & Nolte, 2014). Six equimolar pools (from 35 to 37 individuals per locality) were realized after DNA quality control using Nanodrop, 1,5% agarose gel electrophoresis and Qubit^®^ 2.0 Fluorometer (Invitrogen) quantification.

RAD sequencing was performed following the protocol designed by Etter et al. (2011) and modified by Cruaud et al. (2014). Briefly, DNA pools were digested with 8-cutter restriction enzyme *Sbf*I (21 700 sites predicted following the radcounter_v4.xls spreadsheet available from the UK RAD Sequencing Wiki (www.wiki.ed.ac.uk/display/RADSequencing/Home). For each locality, we built four independent libraries to avoid methodological biases. Digested DNA pools were ligated to a modified Illumina P1 adapter containing locality-specific, 5-6 bp long multiplex identifiers (MIDs). All MIDs differed by at least three nucleotides to limit erroneous sample assignment due to sequencing error (Suppl Mat Table S3). The 24 libraries were then pooled and sheared by sonication using S220 ultra-sonicator (Covaris, Inc.). Genomic libraries were size selected for 300–500 bp by agarose gel excision. P2 adapter were then ligated and fragments containing both adapters (P1 and P2) were PCR enriched during 15 cycles. Libraries were sequenced on an Illumina HiSeq 2000 platform (v3 chemistry) using 2x100 bp paired-end sequencing. Illumina sequencing was performed at the GenePool Genomics Facility (University of Edinburgh, UK).

Sequence reads from Illumina runs were demultiplexed and quality filtered using the *process_radtags* program from the Stacks package version 0.99994. Ambiguous MIDs and low quality reads (Phred < 33) were discarded from further analyses. Sequences were trimmed to 85 nucleotides (position 5 to 90 after the MIDs for the reads 1; position 1 to 85 for the reads 2).

Because no reference genome assembly was available for *M. glareolus*, we needed to build a *de novo* RAD assembly. To that end we first assembled reads 1 per sample with the *ustacks* program from the Stacks package and default options, except for i) the minimum depth of coverage required to create a stack (-m option) that was set to 2; ii) the maximum distance in nucleotides allowed between stacks (-M option) that was set to 3; and iii) the maximum distance allowed to align secondary reads to primary stacks (-N option) that was set to 2. The resulting set loci were then merged into a catalog of loci by the *cstacks* program from the Stacks package run with default options except for the number of mismatches allowed between sample tags to form stacks (-n option) that was set to 2. For each of the obtained read 1 contigs (i.e., RAD loci), we further assembled the associated reads 2 using *CAP3* (Hang & Madan, 1999) ran with default options except for i) the segment pair score cutoff (-i option) that was set to 25; ii) the overlap length cutoff (-o option) that was set to 25; and iii) the overlap similarity score cutoff (-s option) that was set to 400. If a single contig was produced after a first *CAP3* run, this was retained only if it was associated with less than 5% remaining singleton sequences and supported by more than 40 reads. If several contigs were produced, CAP3 was run a second time (using the same options as above) to try to assemble all of them into a single contig which was in this case retained for further analyses. If read 2 contig overlapped with their corresponding read 1 contig, as assessed with the *blastn* program from the BLAST+ v2.2.26 suite (e-value<1e-10 and percentage of identity above 95), both contigs were concatenated. Otherwise, fifteen ‘Ns’ were inserted between both contigs.

Sequence reads were aligned to this assembly using the programs *aln* and sampe implemented in *bwa* 0.5.9 and ran with default options. The resulting *bam* files were then jointly analysed with the *mpileup* program from the Samtools v0.1.19 suite. We used default options except for the minimum mapping quality for alignment (-q option) that was set to 20. The mpileup file was further processed using a custom awk script to perform SNP calling and derive read counts for each alternative base (after discarding bases with a Base Alignment Quality score <25) as previously described (Gautier, 2015; Gautier et al., 2013}. A position was considered variable if (i) it had a coverage of >5 and <500 reads in each pool; (ii) only two different bases were observed across all six pools, and (iii) the minor allele was represented by at least one read in two different pool samples. Note that triallelic positions for which the two most frequent alleles satisfied the above criteria with the third allele represented by only one read were included in the analysis as biallelic SNPs (after filtering the third allele as a sequencing error). To prevent any convergence issue with the SelEstim model-based methods for genome scans that we used (see below), the final dataset was generated by randomizing the reference allele for each and every locus. These procedures were implemented in R (Team 2012) using home-made scripts.

### Genetic variation

#### Diversity at immune-related candidate genes

We performed preliminary analyses on the genotypes inferred at candidate gene SNPs. Observed (*H*o) and expected (*H*e) heterozygosities as well as *F*_IS_ were estimated using GENEPOP v4.2 (Rousset 2008). Deviation from Hardy-Weinberg equilibrium was assessed using exact tests implemented in GENEPOP v4.2. Linkage disequilibrium (LD) was estimated using the program LINKDOS implemented in GENETIX v4.05 (Belkhir, Borsa, Chikhi, Raufaste, & Bonhomme, 1996-2004). Significance was assessed using permutation tests.

#### Characterization of population structure

Population structure analyses were conducted at the population level. We assessed the pattern of differentiation between the six populations pairs (and overall) using the *F*_ST_ estimator developed by Hivert et al. (in prep., script available upon request) for poolseq data. Next, we estimated the scaled covariance matrix of population allele frequencies using the algorithm implemented in BayPass (Gautier, 2015). A principal component analysis (PCA) was performed on this matrix using the FactomineR library in R (Team 2012) to visualize the patterns of population structure. All SNPs were included, as BayPass is likely to be only lightly sensitive to the inclusion of markers evolving under selection when estimating the covariance matrix (Lotterhos & Whitlock, 2014). Finally, we tested for an isolation by distance pattern by analysing the relationship between pairwise genetic distance, estimated as *F*_ST_ / (1 – *F*_ST_), and the logarithm of geographical distance using a Mantel test implemented in R.

### Detecting putative footprints of selection based on differentiation

We used two different methods to characterize markers showing outstanding differentiation (as compared to the rest of the genome) between PUUV endemic and non-endemic areas in Sweden. In these analyses, all candidate loci polymorphisms and RAD SNPs were included.

First, we used the software package SelEstim 1.1.3, which is based on a diffusion approximation for the distribution of allele frequencies in a subdivided population (island model) that explicitly accounts for selection. In particular, SelEstim assumes that each and every locus is targeted by selection to some extent, and estimates the strength of locus-specific selection for each locus, in each subpopulation. SelEstim has been extended since version 1.1.0 to handle Pool-Seq data. Three different SelEstim analyses were run to assess convergence. For each analysis, twenty-five short pilot runs (1 000 iterations each) were set to adjust the proposal distributions for each model parameter and, after a 100 000 burn-in period, 100 000 updating steps were performed. Samples were collected for all the model parameters every 40 steps (thinning interval), yielding 2 500 observations. Convergence was checked using the Gelman–Rubin’s diagnostic implemented in the CODA package for R (Plummer, Best, Cowles, & Vines, 2006). Candidate markers under selection were selected on the basis of the distance between the locus-specific coefficient of selection and a “centering distribution” derived from the distribution of a genome-wide parameter of selection, which accounts for the variation among loci of selection strength. SelEstim uses the Kullback–Leibler divergence (KLD) as a distance between the two distributions, which is calibrated using simulations from a predictive distribution based on the observed data (Vitalis, Gautier, Dawson, & Beaumont, 2014). Hereafter, we report candidate markers with KLD values above the 99.9% quantile of the so-obtained empirical distribution of KLD (although the results based on the 99.95 and 99.99 % quantiles are also provided).

Next, we used the software package BayPass (Gautier, 2015), which extends the approach by Coop et al. (2010) and Günter and Coop (2013). This method relies on the estimation of the (scaled) covariance matrix of population allele frequencies, which is known to be informative about demographic history. Therefore, contrary to SelEstim, BayPass is not limited by the oversimplification of the underlying demographic model. To identify SNPs targeted by selection, we used BayPass to estimate the statistic *X^T^X*, which might be interpreted as a locus-specific analog of *F*_ST_, explicitly corrected for the scaled covariance of population allele frequencies. To define a significance threshold for the *X^T^X* statistic, we used an empirical posterior checking procedure, similar in essence to the one used in SelEstim to calibrate the KLD. The posterior predictive distribution of *X^T^X* was obtained under the null (core) model, by generating and analysing a pseudo-observed dataset (pod) made of 20,000 SNPs (Gautier, 2015). We checked that the scaled covariance matrix of population allele frequencies estimated from the pod was close to the matrix estimated from our data (*FMD* distance = 0.088, see (Gautier, 2015)). The decision criterion for identifying *X^T^X* outliers was defined from the quantiles of the *X^T^X* distribution of the pod analysis.

### Detecting putative footprints of selection based on environmental variables

We used BayPass to test for associations between allele frequencies and environmental variables presumably related to PUUV epidemiology, while controlling for demography. The STD model in BayPass assumes a linear effect of the environmental variable on allele frequencies. We chose empirical Bayesian *P*–values (*eBP*) as the decision criterion, because it was more stable than Bayes Factors (BF) in the sense that estimates were highly correlated across multiple independent runs (see also Bourgeois et al., 2017). Similarly, MCMC sample based estimate of BF (-auxmodel option) or eBP (-covmcmc option) were inaccurate likely due to identifiability issues related to the too small number of populations considered. Roughly speaking, for a given SNP, the empirical Bayesian *P*–value measures to which extent the posterior distribution of the regression coefficient excludes 0 (Gautier, 2015). In order to compute the *eBP*, we used the importance sampling algorithm implemented in BayPass to estimate the moments of the posterior distribution of the regression coefficients. We calibrated *eBP* using simulations from a posterior predictive distribution, based on the observed data set.

Environmental variables related to PUUV prevalence in human (number of nephropathia epidemica cases), climate, forest composition and shape, and soil water content, were selected with regard to PUUV ecology in Europe as they should reflect PUUV distribution in Sweden (Zeimes et al., 2012; Zeimes et al., 2015) (Table 2). Except for the number of nephropathia epidemica cases that was only available per county, variables were computed within an area covering a circular radius of 3 km around each sampling site (ArcGIS 10.1), which is an acceptable estimate of vole dispersal capacity (Le Galliard, Rémy, Ims, & Lambin, 2012). To summarize climate variation, we used the minimum temperature in winter (December, January and February), the maximum temperature in summer (June, July and August), the percentage of the area covered by snow and the annual precipitation. Land-use was characterized by forest types (the proportion in the 3km buffer of forest, coniferous, broadleaved and mixed forest) and by tree species (the volume of spruce and pine and their standard deviation). Forest patches metrics (computed with FRAGSTATS, version 4, http://www.umass.edu/landeco/research/fragstats/fragstats.html) were averaged in the 3km buffer and included the contiguity index, the shape index (a shape index of one represents the most compact shape, upper than one, a more complex shape) and the perimeter. Finally, the soil water index (SWI) representing the soil moisture conditions was also included.

**Table 2.** Environmental parameters and their potential impacts (indicated by ‘x’) on the bank voles’ abundance or virus survival outside the host (see Zeimes et al., 2015 for references).

| Variables | Bank voles | Virus | Resolution | Units | Sources |
| --- | --- | --- | --- | --- | --- |
| PUUV presence |  |  |  |  |  |
| Total number of human cases between<br>2001 and 2011 (SWI dataset) |  | x | County |  |  |
| Climatic variables |  |  |  |  |  |
| Minimum temperature in winter<br>(MinTempWinter) | x | x | 0.0083°<br>1950-2000 | °C | Worldclim |
| Maximum temperature in summer<br>(MaxTempSummer) |  | x | 0.0083°<br>1950-2000 | °C | Worldclim |
| Snow cover (Snow) | x | x | 0.005°<br>2000-2008 | Area percentage | MODIS |
| Annul precipitation (Pp) | x | x | 0.0083° | mm | Worldclim |
|  |  |  | 1950-2000 |  |  |
| Forest and soil indices |  |  |  |  |  |
| Proportion of forest | x |  | 100 m | Area percentage | Corine 2006 (EEA) |
| Proportion of broadleaved forest | x |  | 100 m | Area percentage | Corine 2006 (EEA) |
| Proportion of coniferous forest | x |  | 100 m | Area percentage | Corine 2006 (EEA) |
| Proportion of mixed forest | x |  | 100 m | Area percentage | Corine 2006 (EEA) |
| Mean volume of spruce | x |  | 25 m | m<sup>3</sup>/ha | SLU Skogskarta |
| Standard deviation of the volume of spruce | x |  | 25 m | m<sup>3</sup>/ha | SLU Skogskarta |
| Mean volume of pine | x |  | 25 m | m<sup>3</sup>/ha | SLU Skogskarta |
| Standard deviation of the volume of pine | x |  | 25 m | m<sup>3</sup>/ha | SLU Skogskarta |
| Average contiguity index of forest patches | x |  | 1:20000 | Relative unit | Lantmäteriet |
| Average shape index of forest patches | x |  | 1:20000 | Relative unit | Lantmäteriet |
| Perimeter of the forest patches | x |  | 1:20000 | m | Lantmäteriet |
| Soil water index (SWI) | x | x | 25 km<br>2007-2010 | Relative unit | TU-WIEN |

To reduce the dimensionality of these environmental data, we assessed SNP-environment associations using the two first principal components (PC) from a PCA that included all 15 variables (Fig. 2). These new synthetic variables explained respectively 44.1% and 36.6% of the total variance. PC1 represented an environmental, latitudinal gradient. Along this axis, sampling localities were ranked from northern localities (positive values) exhibiting high numbers of human PUUV infection and large volume of spruce forests, to southern localities (negative values) with mixed or broadleaved forests, low mean winter and high maximum summer temperatures. PC2 strongly opposed the two northern sampling localities, with Hörnefors (negative values) being characterized by large volume of contiguous coniferous forest and high snow coverage, and Harnosand (positive values) being more fragmented.

**Fig. 2.**
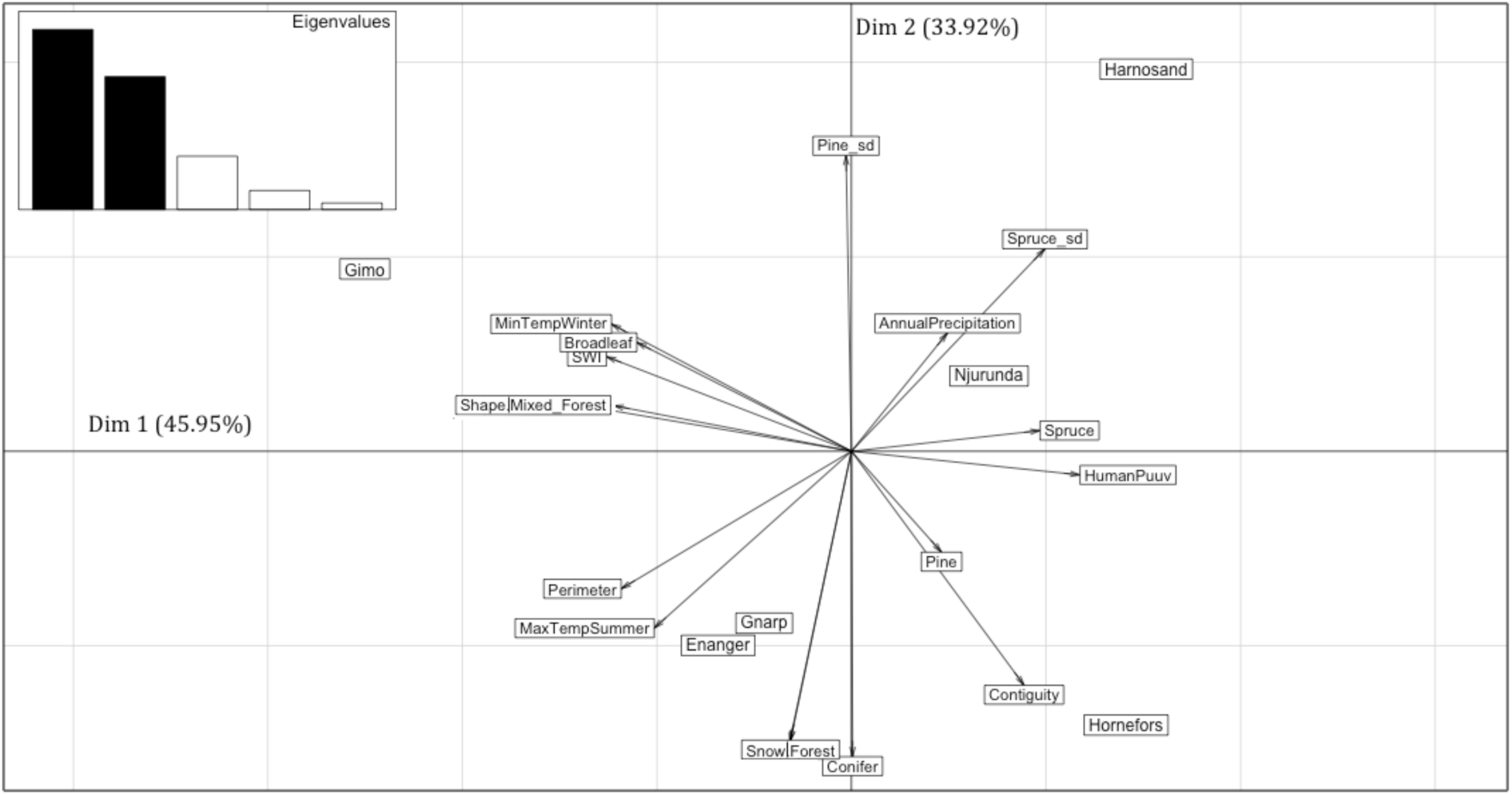
Principal Component Analysis (PCA) plot of the environmental data and of the six populations analysed. Details about envionmental variables are provided in Table 2.

### Annotations and gene ontology analysis

#### Annotation of contig sequences

We applied several approaches to functionally annotate the contigs. First, all consensus sequences (reads1-reads2) were blasted against the NCBI NT (v. 03/29/2015) and NR (v. 05/16/2015) databases using the blastn (v. 2.2.28+, parameters: threshold e-value of 1e-5, minimum alignment percent identity of 70%) and blastx (e-value of 1e-5) search algorithms, respectively. In-house Perl scripts were applied to further filter these BLAST results. Only matching sequences of the taxon ‘*rodentia*’ were considered.

In addition, in the absence of a published complete genome of *M. glareolus*, we compared the RAD contigs to the genome of *Mus musculus* (build GRCm38/mm10). We downloaded (date: 2015-06-25) the sequences of the 22 mouse chromosomes, 90,891 cDNAs and 87,139 proteins from ENSEMBL (http://www.ensembl.org/) and blasted the RAD contigs against these sequences (blastn for genome and cDNA, blastx for proteins with same parameters as mentioned above).

#### Gene enrichment analysis of outlier candidates

For enrichment analyses of metabolic pathways and Gene Ontologies (GO), we applied the KOBAS web-application (KOBAS 2.0, http://kobas.cbi.pku.edu.cn). For this purpose, we extracted the ENSEMBL gene ids from the blastx results of the RAD outliers versus the ENSEMBL protein dataset and also included the gene id of the candidate *Tlr7*. The metabolic pathway databases KEGG, Reactome, BioCyc and PANTHER, as well as the GO database were chosen for the enrichment analysis. The genome of *Mus musculus* (GRCm38) was chosen as species of interest. On both ENSEMBL id sets the ‘annotate’ and ‘identify’ programs of KOBAS were executed with the ENSEMBL ids of the unique set of outliers (all methods combined) as sample file and the ids of the entire GRCm38 coding gene set as background dataset. For these analyses, the other parameters of the ‘identify’ program were left on default settings. Therefore, the enrichment analysis was done using the hypergeometric and Fisher’s exact tests, and Benjamini and Hochberg’s method was applied for FDR correction (Benjamini & Hochberg, 1995). A metabolic pathway was considered as significantly enriched, if the associated *q*-value was less than or equal to 0.05. Finally, a full network representation was provided using the Search Tool for the Retrieval of Interacting Genes (STRING) database (Snel, Lehmann, Bork, & Huynen, 2000). Furthermore, we ran the web program REVIGO (web version of 07/27/2015, http://revigo.irb.hr/; http://www.ncbi.nlm.nih.gov/pmc/articles/PMC3138752/) to group significantly enriched Gene Ontology (GO) terms into GO categories and to summarise the GO terms to common terms. We applied the SimRel semantic similarity measure with the default GO term similarity, associated *q*-values to the enriched GO terms, and the *Mus musculus* database (Supek, Bošnjak, Škunca, & Šmuc, 2011).

## Results

### RAD tag sequencing

Sequencing of the 24 RAD libraries (6 localities and 4 replicates) generated 340,692,418 reads, with an average of ca. 13.5 million reads per Multiplex Identifiers (MIDs). The number of sequences generated per locality and MID ranged between ca. 10.1 and ca. 15.1 million reads. After trimming sequences to 85 bp and after filtering for quality, an average of ca. 12.6 million reads per MID (representing 93.5% of the total) were retained. Read 1 assembling produced 151.522 contigs. The first *CAP3* assembling run of the associated reads 2 produced 46474 unique contigs represented by more than 40 sequences, and among which 46.471 had less than 5% singleton. The second *CAP3* assembling run enabled to provide 69.777 read 2 contigs. We found a single significant alignment between read 1 and read 2 contigs in 59.242 cases and no significant overlap in 10.495 cases. The other 38 cases corresponded to a complete alignment of read 1 contig with read 2 contig (1 case), an alignment of read 1 contig within read 2 contig (2 cases) and multiple significant alignment (35 cases). The resulting assembly finally consisted of 69,777 contigs spanning 38.482 Mb (average contig size equal to 551.5, [209-891]).

Reads were aligned to this reference contig dataset and 485,182 SNPs were detected (QC > 25, depth range: 5-500X). Among them 95,988 SNPs distributed on 70,699 contigs were kept according to the more stringent criteria described before.

### Descriptive statistics

#### Candidate gene diversity

The Sanger sequencing of the four immunity-related genes for 12 individuals identified a total of 20 variable sites (19 SNPs and one insertion-deletion event) from 5395 bp sequence data (Suppl. Mat. Table S4). Each gene had between one (*Tlr7*) and 14 (*Tlr4*) SNPs. Eight of these SNPs were polymorph in Sweden and were successfully genotyped in 250 bank voles using the KASP genotyping (Table 3).

**Table 3.**
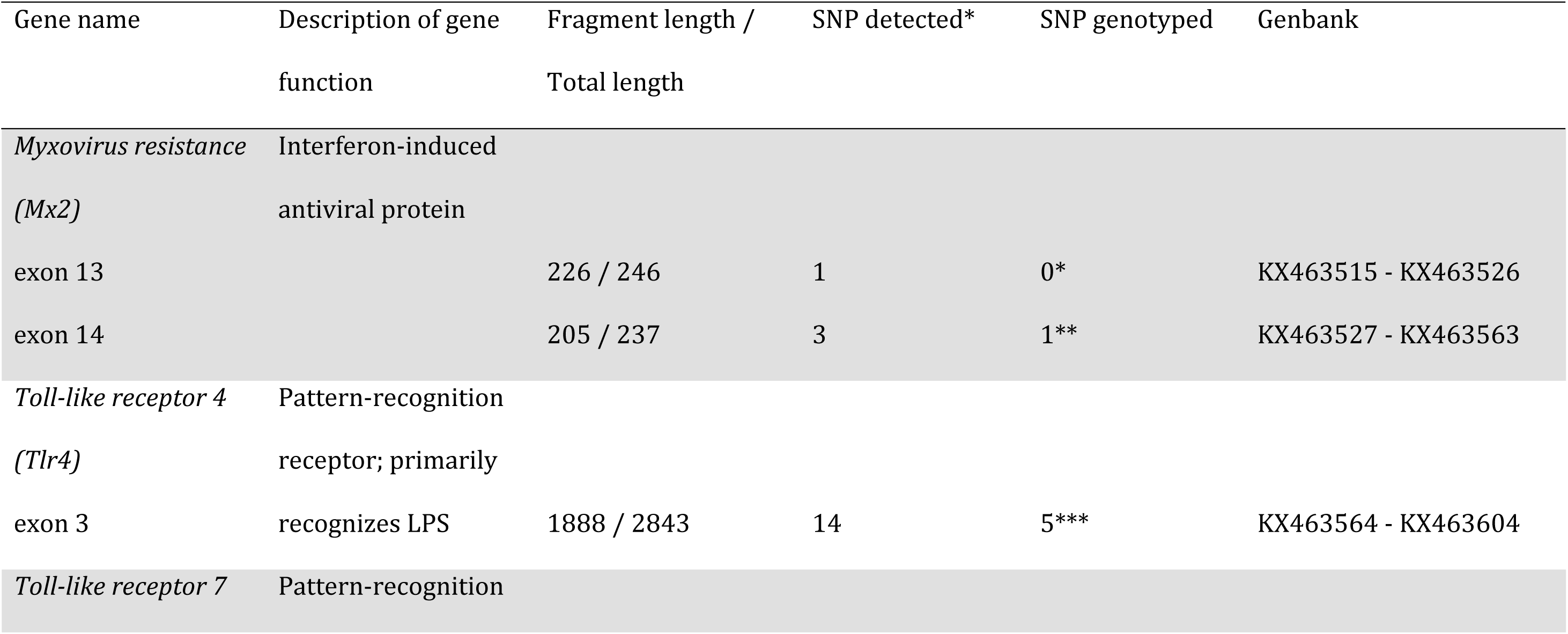
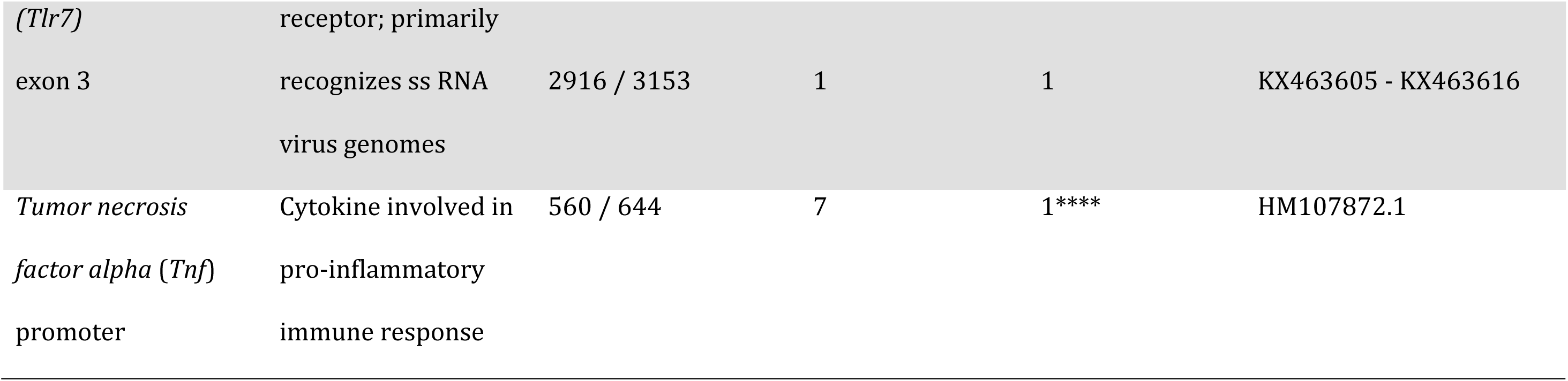
Summary of sequenced candidate immune-related genes of bank voles (n=12). More details about these SNPs are provided in Table S3. * This insertion/deletion was only polymorph in Germany. ** one was not genotyped because of 100% LD with the other one, the non synonymous SNP was only polymorph in France. *** The other SNPs were either only detected in one individual from France or only detected in Germany. **** Only one SNP shown to be involved in PUUV / *M. glareolus* interaction (Guivier et al., 2014; Guivier et al., 2010) was genotyped.

Not surprisingly, significant linkage disequilibrium (LD) was observed between most SNPs located within genes: Tlr4-exon3 776 and Tlr4-exon3 1146, Tlr4-exon3 1662, Tlr4-exon3 1687; Tlr4-exon3 1146 and Tlr4-exon3 1662, Tlr4-exon3 1687; Tlr4-exon3 1662 and Tlr4-exon3 1687. We did not detect any linkage disequilibrium among SNPs located in different genes.

Estimates of diversity indices per SNP and per sampling locality can be found in Table 4. Deviation from Hardy-Weinberg equilibrium was observed in most localities for *Tnf* promoter (-296) with significant deficits in heterozygotes detected in Hörnesand, Harnefors, Njurunda and Gimo. Moreover, significant departures from Hardy-Weinberg expectations were observed in Gimo for all polymorphic SNPs in this locality.

**Table 4.** Diversity indices per sampling locality and SNP. Values in bold indicate significant *p*-values.

| SNP | Indices | Localities |  |  |  |  |  |
| --- | --- | --- | --- | --- | --- | --- | --- |
|  |  | Hörnesand | Härneförs | Njurunda | Gnarp | Enånger | Gimo |
| Mx2_14_162 | He n.b. | 0.4639 | 0.1759 | 0.000 | 0.000 | 0.000 | 0.000 |
|  | Hobs. | 0.4694 | 0.1579 | 0.000 | 0.000 | 0.000 | 0.000 |
|  | Fis W.C. | -0.012 | 0.103 | - | - | - | - |
| TLR4_667 | He n.b. | 0.000 | 0.000 | 0.0294 | 0.1311 | 0.1726 | 0.5065 |
|  | Hobs. | 0.000 | 0.000 | 0.0294 | 0.1389 | 0.1875 | 0.2703 |
|  | Fis W.C. | - | - | 0.000 | -0.061 | -0.088 | **0.470** |
| TLR4_776 | He n.b. | 0.0600 | 0.0517 | 0.2950 | 0.4261 | 0.4583 | 0.2755 |
|  | Hobs. | 0.0612 | 0.0526 | 0.2941 | 0.4286 | 0.5000 | 0.1622 |
|  | Fis W.C. | -0.021 | -0.018 | 0.003 | -0.006 | -0.093 | **0.415** |
| TLR4_1146 | He n.b. | 0.0791 | 0.0517 | 0.2950 | 0.4409 | 0.4583 | 0.2755 |
|  | Hobs. | 0.0816 | 0.0526 | 0.2941 | 0.4167 | 0.5000 | 0.1622 |
|  | Fis W.C. | -0.032 | -0.018 | 0.003 | 0.056 | -0.093 | **0.415** |
| TLR4_1662 | He n.b. | 0.0600 | 0.0517 | 0.3021 | 0.4343 | 0.4583 | 0.2936 |
|  | Hobs. | 0.0612 | 0.0526 | 0.3030 | 0.4054 | 0.5000 | 0.1892 |
|  | Fis W.C. | -0.021 | -0.018 | -0.003 | 0.067 | -0.093 | **0.359** |
| TLR4_1687 | He n.b. | 0.0412 | 0.0517 | 0.2950 | 0.4409 | 0.4583 | 0.2755 |
|  | Hobs. | 0.0417 | 0.0526 | 0.2941 | 0.4167 | 0.5000 | 0.1622 |
|  | Fis W.C. | -0.011 | -0.018 | 0.003 | 0.056 | -0.093 | **0.415** |
| TLR7_2593 | He n.b. | 0.0000 | 0.3552 | 0.1383 | 0.4261 | 0.5079 | 0.0000 |
|  | Hobs. | 0.0000 | 0.1404 | 0.1471 | 0.3143 | 0.3750 | 0.0000 |
|  | Fis W.C. | - | **0.607** | -0.065 | 0.265 | 0.265 | - |
| TNFp_296 | He n.b. | 0.4033 | 0.4988 | 0.3652 | 0.2950 | 0.2285 | 0.4832 |
|  | Hobs. | 0.0204 | 0.0175 | 0.1765 | 0.2941 | 0.1935 | 0.2432 |
|  | Fis W.C. | **0.950** | **0.965** | **0.521** | 0.003 | 0.155 | **0.500** |

#### Characterization of population structure

The multilocus *F*_ST_ between pairs of populations ranged from 0.091 to 0.361, and the overall differentiation among populations was estimated as *F*_ST_ = 0.212 (Suppl. Mat. Fig. S1). We found a significantly positive correlation between pairwise genetic distance, estimated as *F*_ST_ / (1 - *F*_ST_), and the logarithm of geographical distance (Mantel test; *p* = 0.0014). The PCA performed on the covariance matrix of population allele frequencies revealed a strong differentiation between the northern populations Hörnefors (mitochondrial lineage ‘Ural’), Härnösand (mitochondrial lineage ‘western’) and all other populations on the first axis. The second axis differentiated the southern population of Gimo from all other populations (Fig. 3a). The scaled covariance matrix of population allele frequencies estimated with BayPass was also consistent with a strong differentiation between Hörnefors and Härnösand populations on the one hand, and more southern populations on the other hand (Fig. 3b), as well as with an isolation-by-distance pattern (Fig. 3c).

**Fig. 3.**
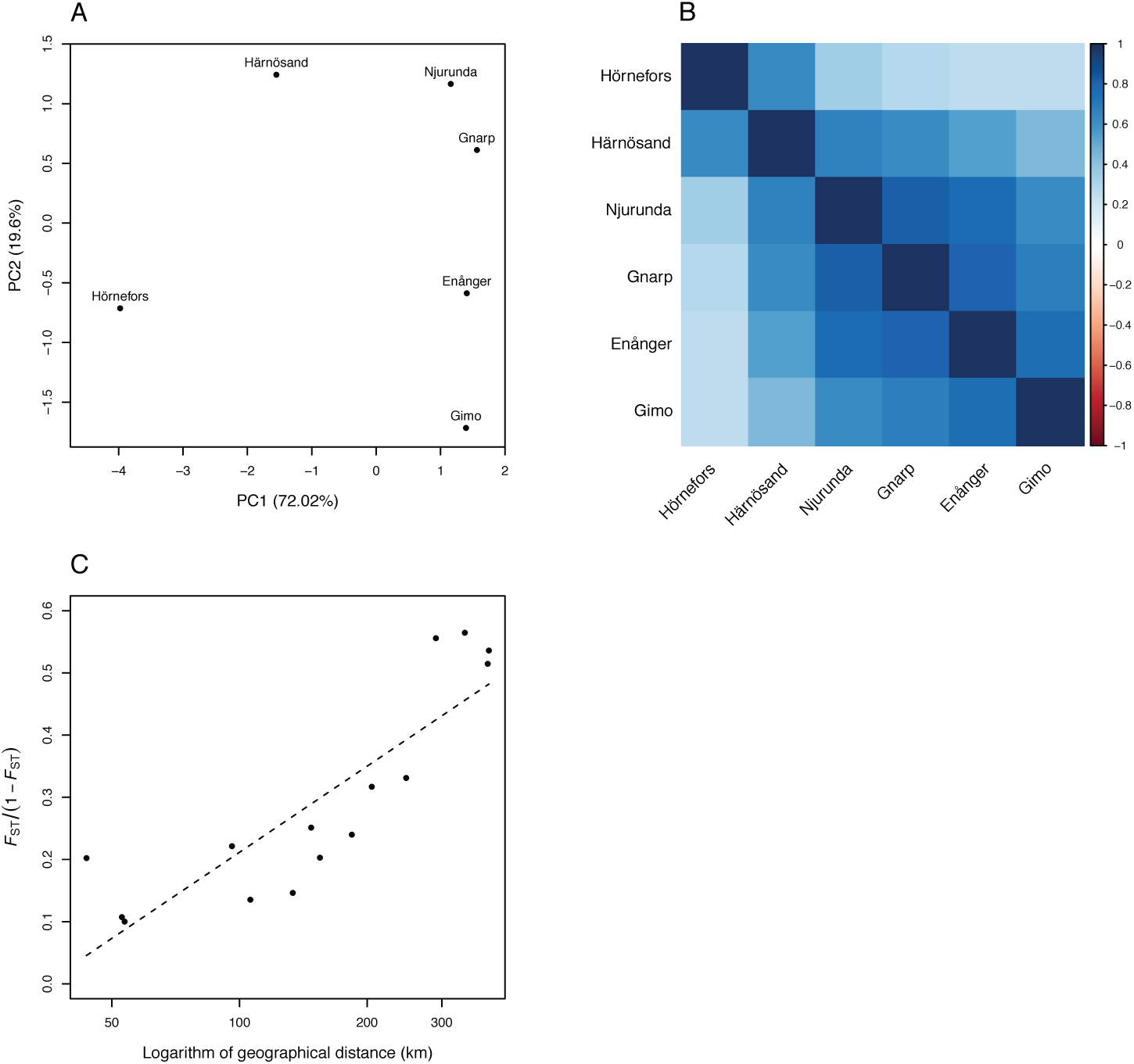
Graphical representations of the *M. glareolus* population genetic structure based on the 95,988 SNPS. a) Principal Component Analysis (PCA) based on the variance covariance matrix of the six bank vole populations studied, estimated using BayPass and based on the 95,988 SNPs included in the statistical analyses. b) Representation of the scaled covariance matrice as estimated from BayPass under the core model with *p* = 1. c) Isolation by distance pattern.

### Signatures of selection

#### SelEstim

The Gelman–Rubin’s diagnostic was equal to 1.06 for the hyper-parameter *λ*, which represents the genome-wide effect of selection over all demes and loci, and to 1.11 for the parameters *M*, which represent the scaled migration parameters. This indicates that the chains converge satisfactorily to the target distribution. One replicate analysis was therefore picked at random for the rest of the study. The 99.9% quantile of the posterior predictive distribution of the KLD (based on pseudo-observed data) equalled 2.61. We found a total of 86 SNPs, representing 78 unique contigs and the candidate gene *Tlr7*, with a KLD estimate equal to or larger than this threshold (48 SNPs were identified as outliers using the 99.95% quantile threshold, and 37 at the 99.99% threshold). All these outliers showed high *F*_ST_ estimates compared with the background *F*_ST_ (Fig. 4a). We found a clinal pattern of variation for the locus- and population-specific coefficients of selection estimated by SelEstim along the North / South axis of sampling. Furthermore, we found, for most outliers, that the coefficients of selection were correlated with the first principal components of the environmental variables that discriminate areas of high PUUV prevalence in humans from non-endemic PUUV areas (Fig. 4b).

**Fig. 4.**
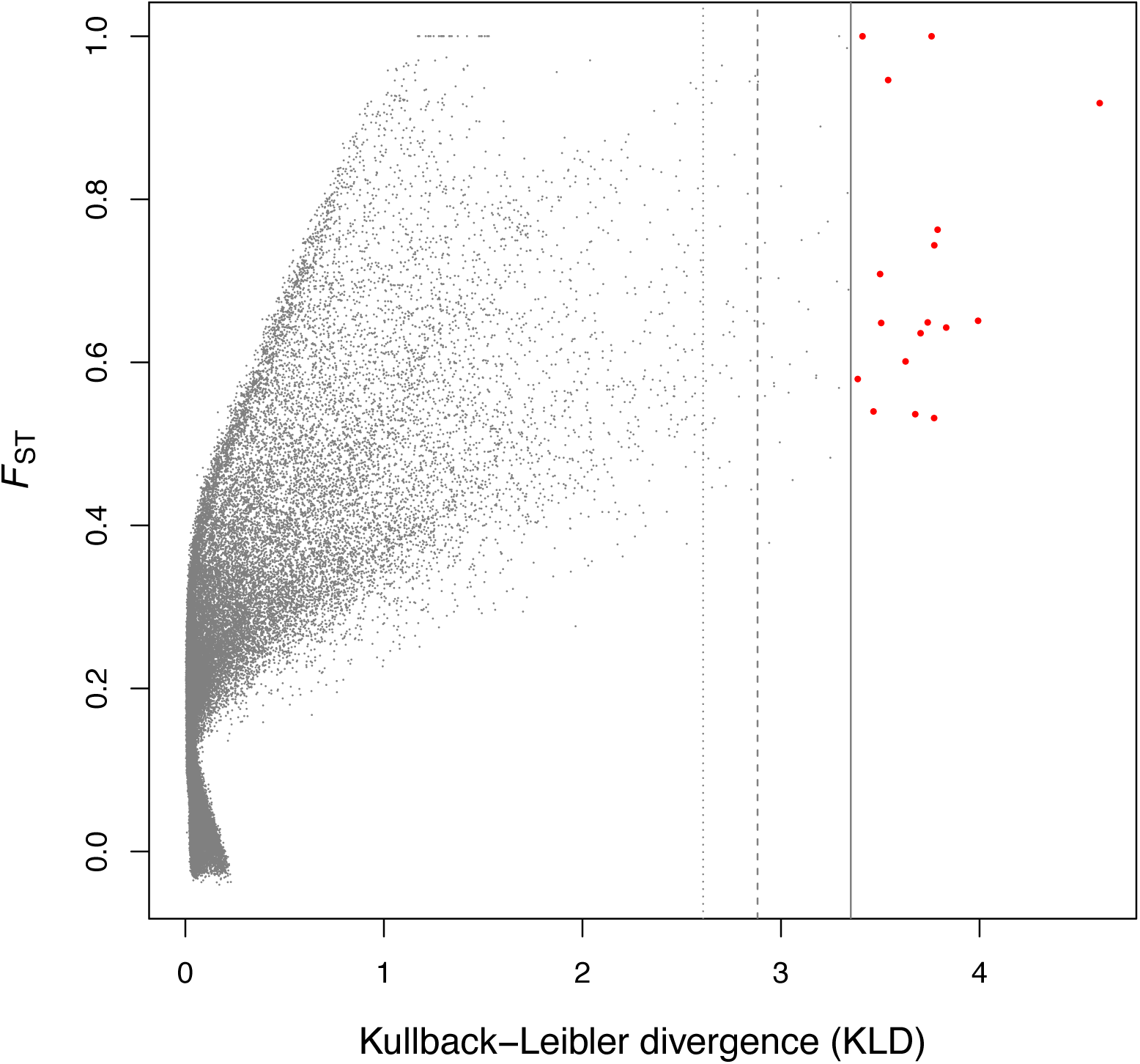
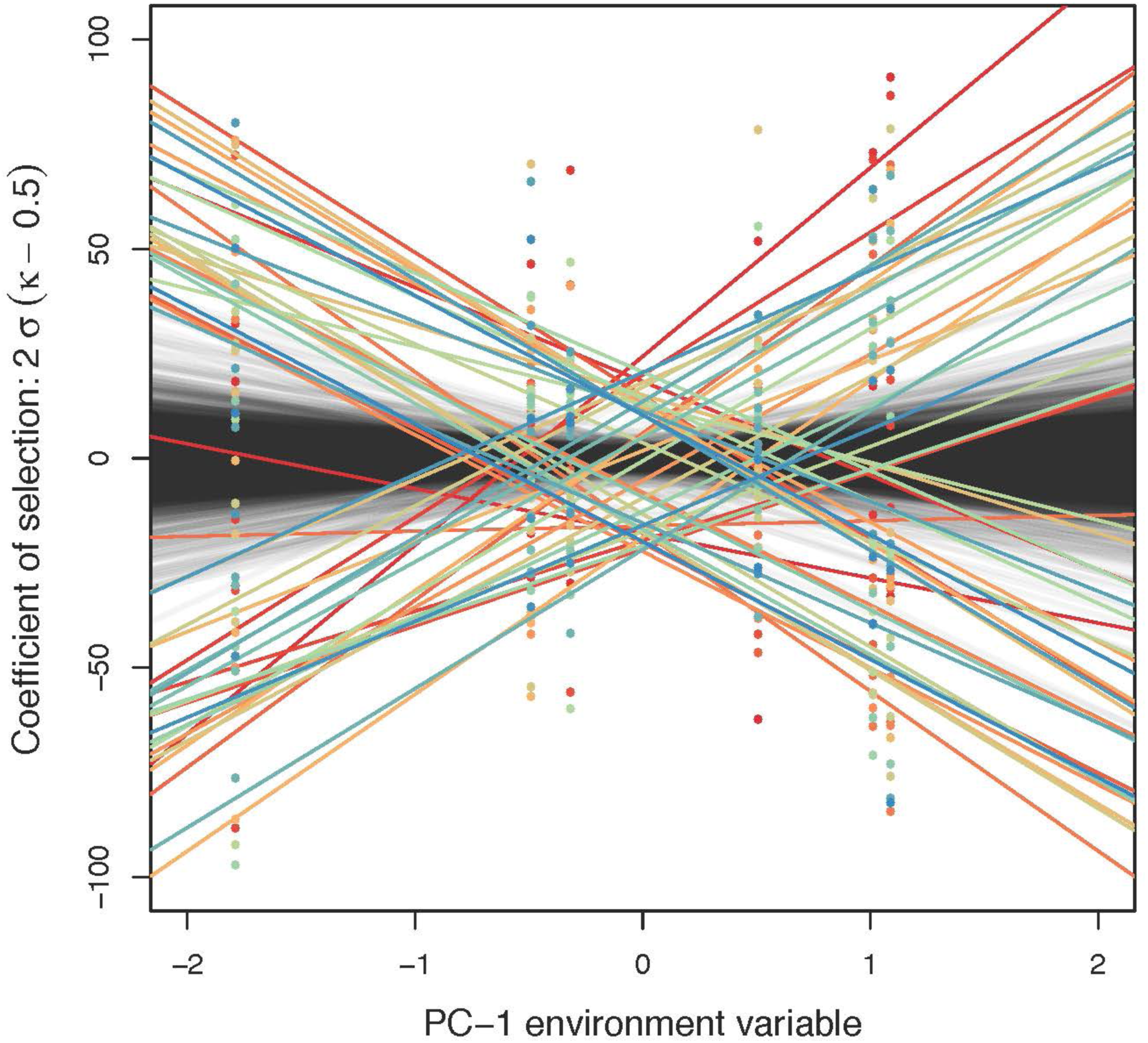
Outlier detection based on 95,988 SNPs in the six bank vole populations using SelEstim. a) *F*_ST_ estimates are represented as a function of the Kullback-Leibler Divergence (KLD) measure for all SNPs. Vertical lines correspond to the 99.90, 99.95 and 99.99 % quantiles calculated from the KLD calibration procedure. b) Correlations between the locus-and population-specific coefficient of selection and the coordinates of the first principal components of the environmental variables. The colored lines stand for outlier SNPs, and the thin grey lines stand for all markers. The coefficient of selection was transformed as: 2 * \sigma_{ij} * (\kappa_{ij} – 0.5).

#### BayPass

Considering the core model implemented in BayPass (i.e., the covariable-free approach), we found 10 outlier SNPs, belonging to nine unique contigs (*X^T^X* > 14.88, 99.9% quantile). Six SNPs corresponding to five unique contigs were common with the SelEstim analysis (Fig. 5a). Considering the 99.95% and the 99.99% quantiles we found five and one outlier SNP(s), respectively. Out of these, three (respectively zero) were common with the SelEstim analysis (Suppl. Mat. Fig. S2).

**Fig. 5.**
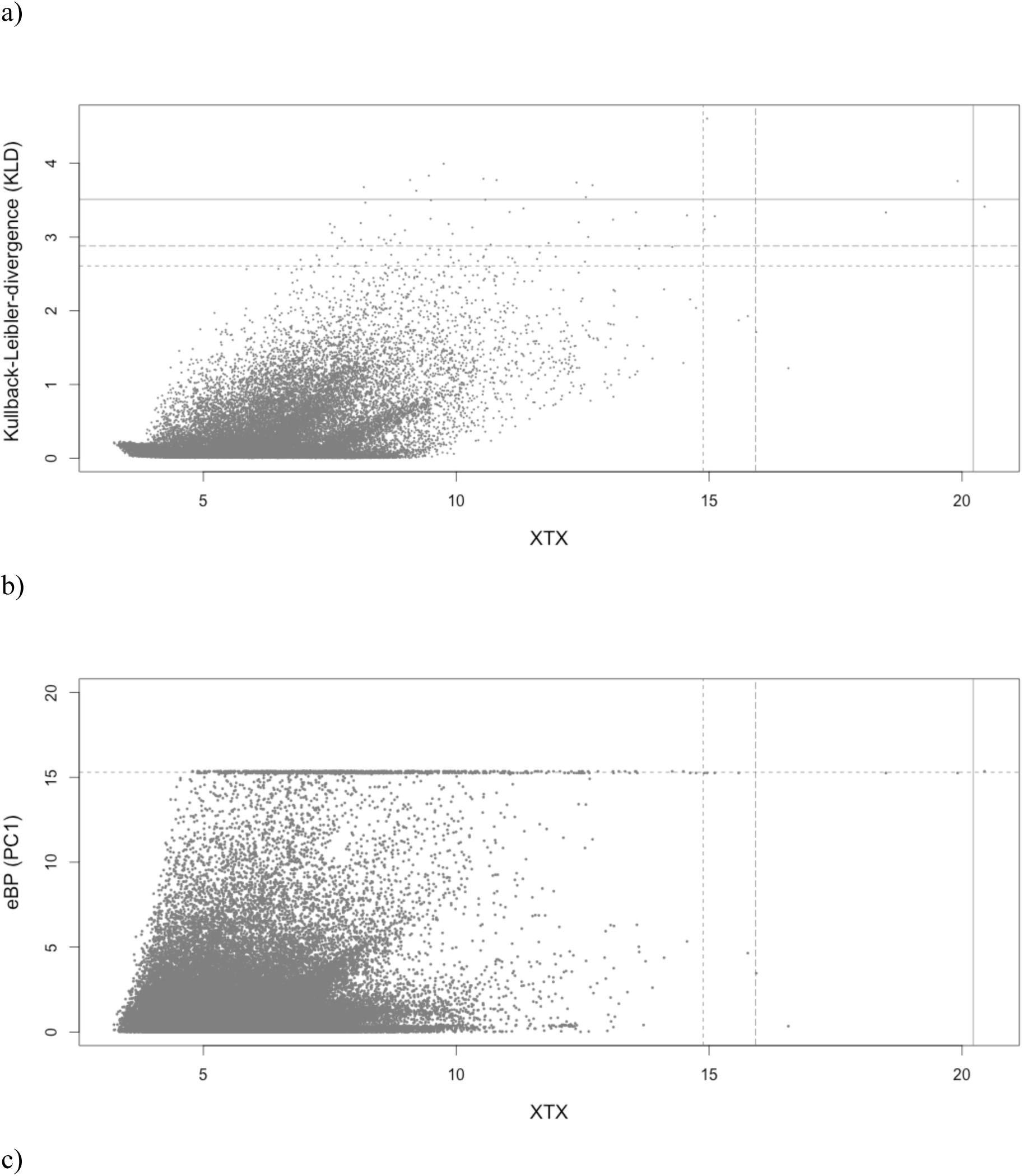
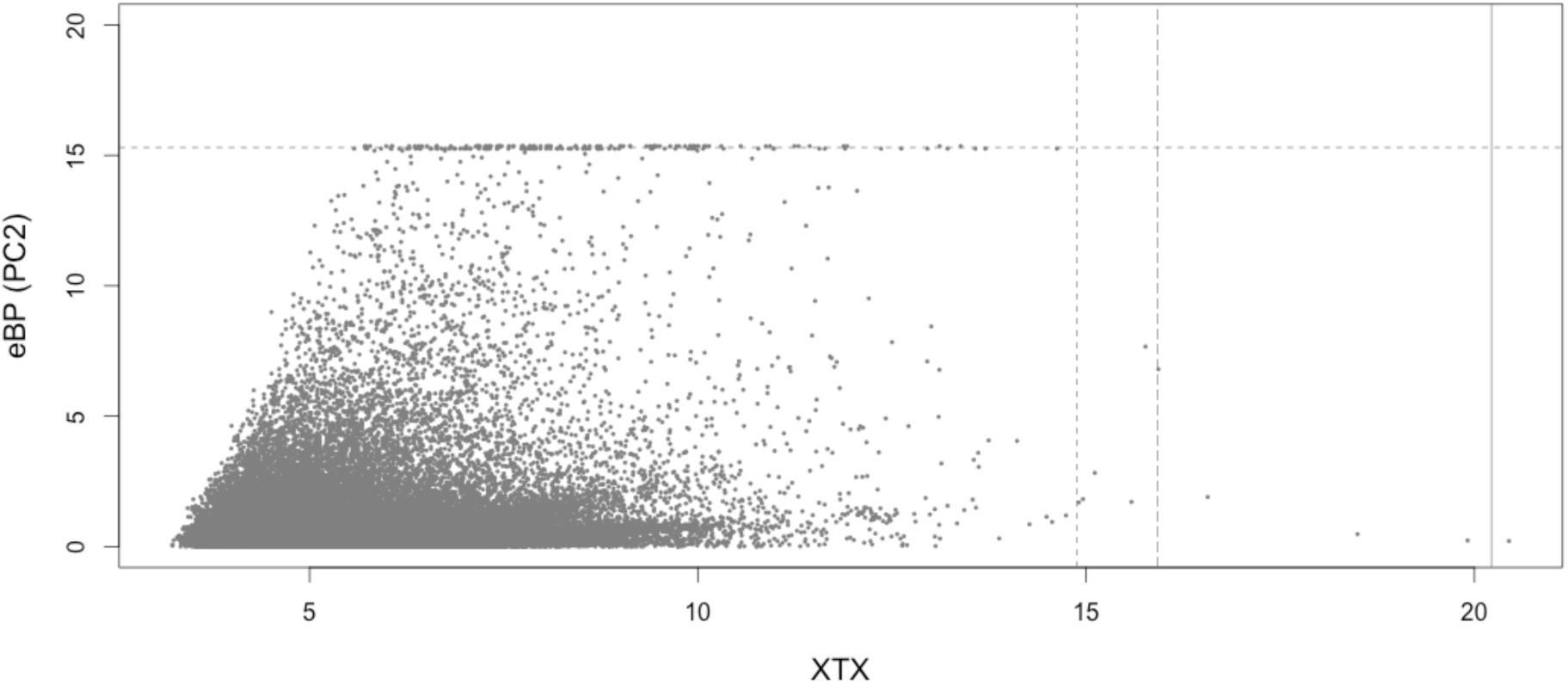
Outlier detection based on 95,988 SNPs in the six bank vole populations using BayPass. a) Correlation between Kullbac-Leibler Divergence (KLD) measure for all SNPs and the statistics X^T^X estimated using BayPass. Horizontal lines correspond to the 99.90, 99.95 and 99.99 % quantiles calculated from the KLD calibration procedure. Vertical lines correspond to these three quantiles calculated using simulations from a predictive distribution based on the observed dataset. b) Correlation between the statistics *eBP* (with the environmental synthethic variable being PCA axis 1) and X^T^X estimated using BayPass. Horizontal and vertical lines correspond to the 99.90, 99.95 and 99.99 % quantiles calculated from the simulations described above. c) Correlation between the statistics *eBP* (with the environmental synthethic variable being PCA axis 2) and X^T^X estimated using BayPass. Horizontal and vertical lines correspond to the 99.90, 99.95 and 99.99 % quantiles calculated from the simulations described above.

Using the STD model in BayPass (which allows the evaluation of associations between SNP allele frequencies and environmental variables), we found 483 SNPs (belonging to 413 unique contigs) with strong association signals (*eBP* > 15.35). A total of 395 SNPs – corresponding to 339 unique contigs – showed significant association with the first principal component of the environmental variables, that discriminate areas of high PUUV prevalence in humans from non-endemic PUUV areas (Fig. 5b). Among them, 26 contigs were previously detected using SelEstim only (24 contigs) or SelEstim and BayPass core model (2 contigs). Ninety SNPs – corresponding to 77 unique contigs – were associated with the second principal component of the environmental variables (Fig. 5c). Four of these contigs were previously detected as outliers: three from the BayPass STD model with the first PC, and another one from the SelEstim analysis. None of these SNPs were found as outliers in the BayPass core model. Results obtained with other threshold values are provided in Suppl. Mat. Fig. S2.

### Annotation and gene ontology analysis

#### Annotation of contig sequences

Blastn provided an annotation for 49.6% (34,578 contigs) of 69,777 bank vole contigs tested across Rodentia databases; blastx led to an annotation for 19.8% (13,830 contigs) of them. In total, the BLAST similarity matches could be assigned to twelve rodent species. The Chinese hamster (37.5%), the house mouse (27.3%) and the Norway rat (17%) were among the most represented rodents. When considering the genome sequence of *M. musculus* obtained from ENSEMBL, we observed that all 22 mouse chromosomes were covered by RAD contigs (*n* = 30,929, i.e., 44.3%). Moreover, about 20% (*n* = 13,856) of the 69,777 bank vole contigs were located in 6,077 mouse protein-coding genes (cDNAs). Out of these, 9,606 (13.8%) bank vole RAD contigs were actually located in the coding sequence of 4,706 mouse proteins (the other RAD contigs were more likely located in UTR termini of the cDNA).

#### Annotation, function and gene enrichment analysis of outlier candidate genes

Gene enrichment analysis was performed on the 468 bank vole outlier contigs detected by at least one of signature selection methods. Among them, 191 were aligned to the mouse genome, covering 20 mouse chromosomes. In total, 52 outliers – including the *Tlr7* candidate gene – matched to 44 mouse protein-coding genes (Table 5). Note that 10 of these genes were coding for proteins that are involved in immunity (*Dgkd*, *Fermt3, Il12rb1, Lbp, Lilrb4, Nedd4, Ptprc, Tlr7, Tnfrsf22, Vwa*). They were either detected by the algorithms of SelEstim or BayPass with environmental associations.

**Table 5.**
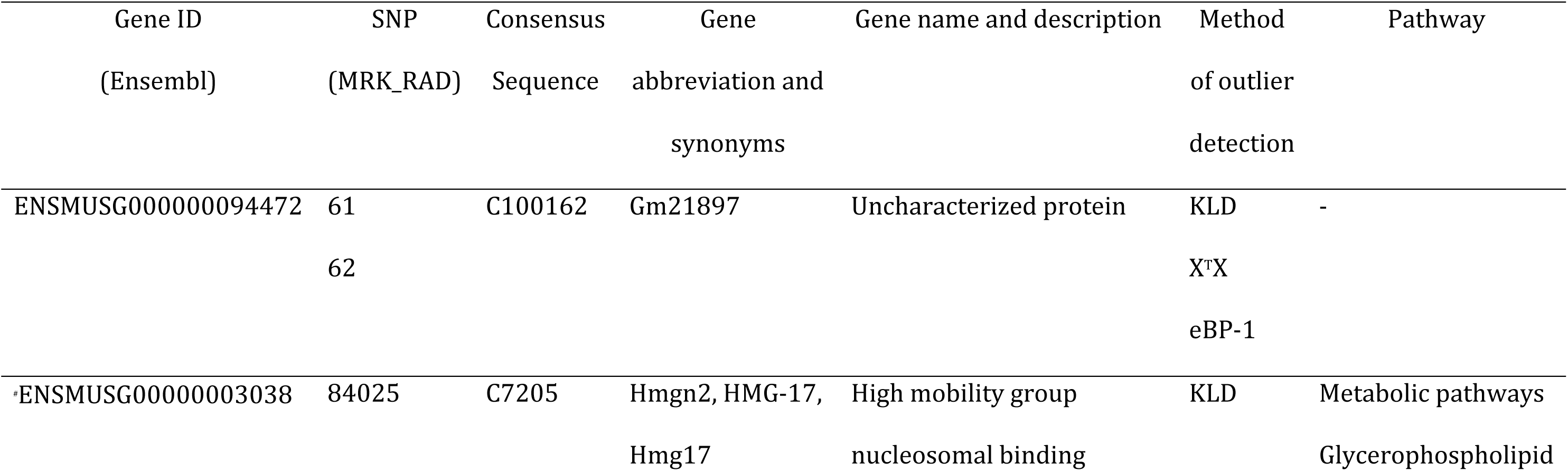
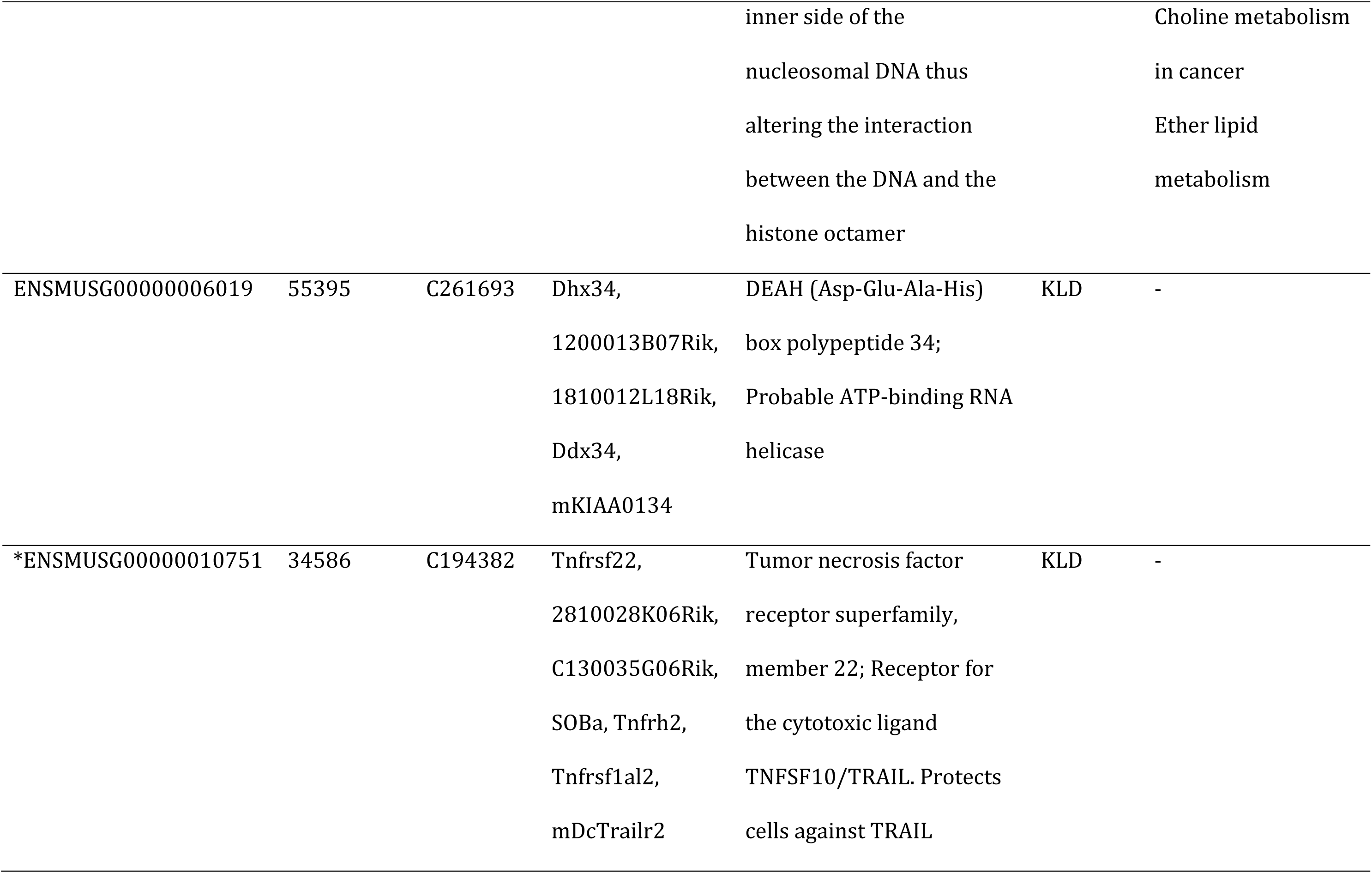
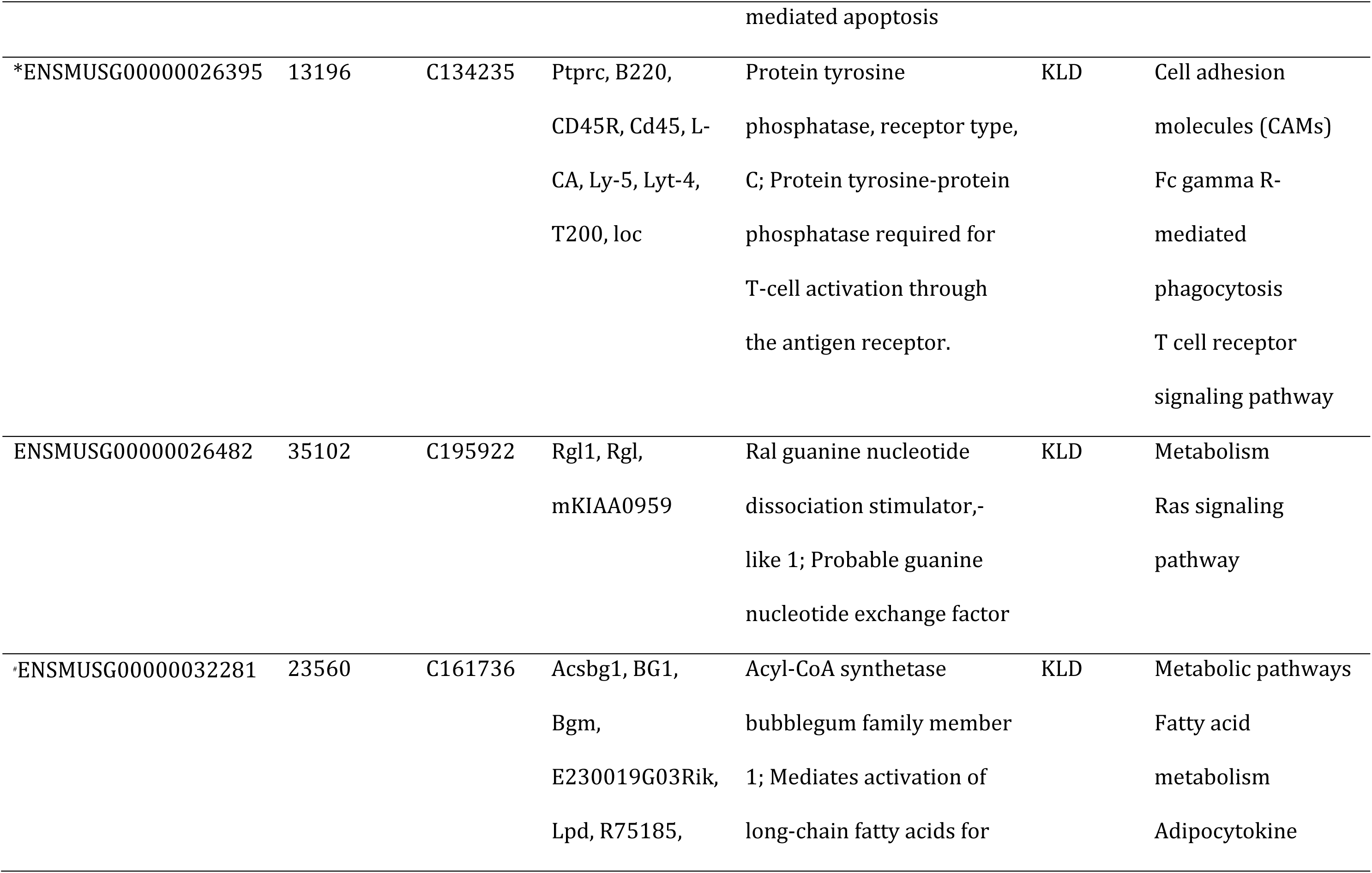
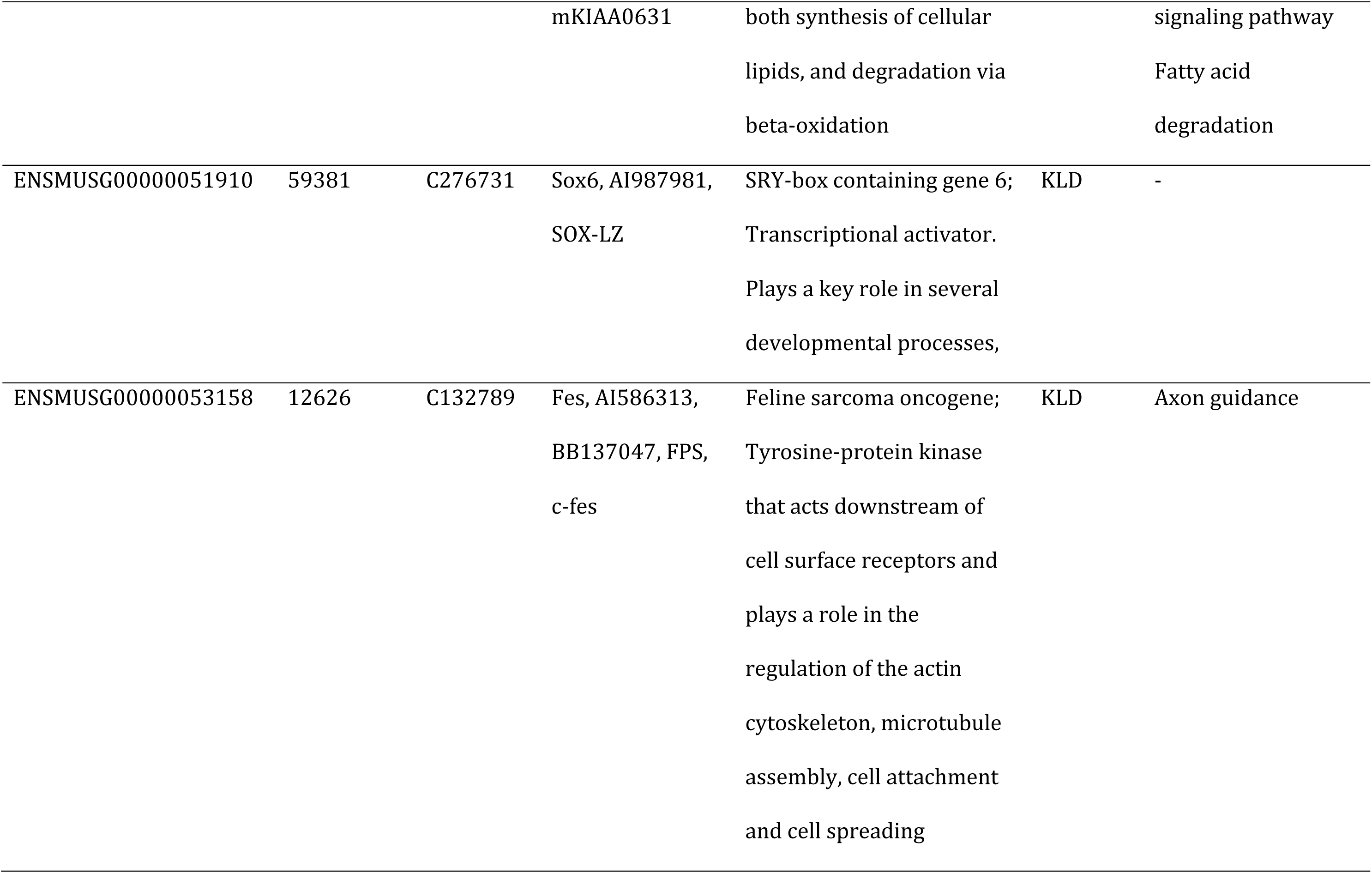
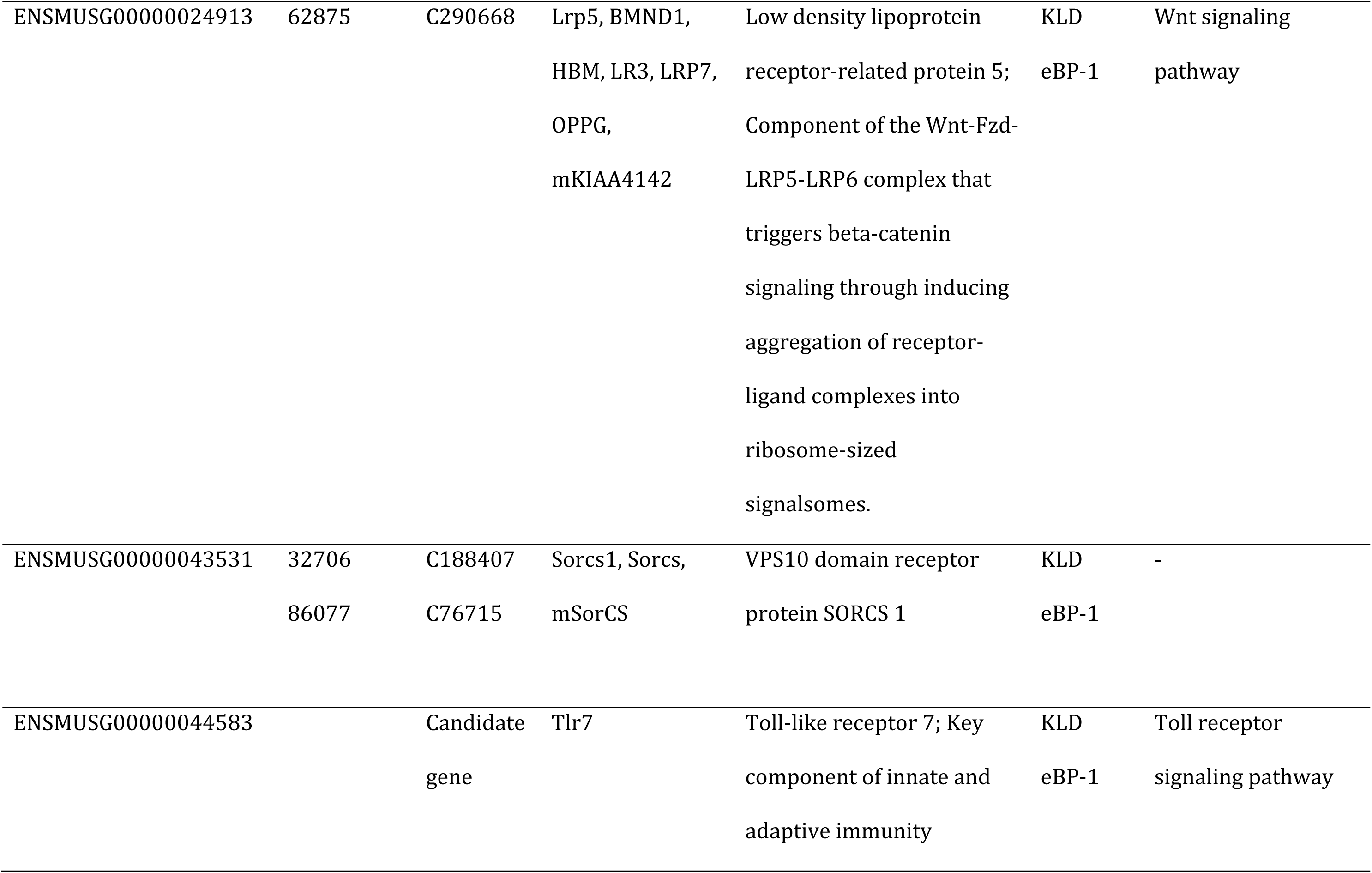
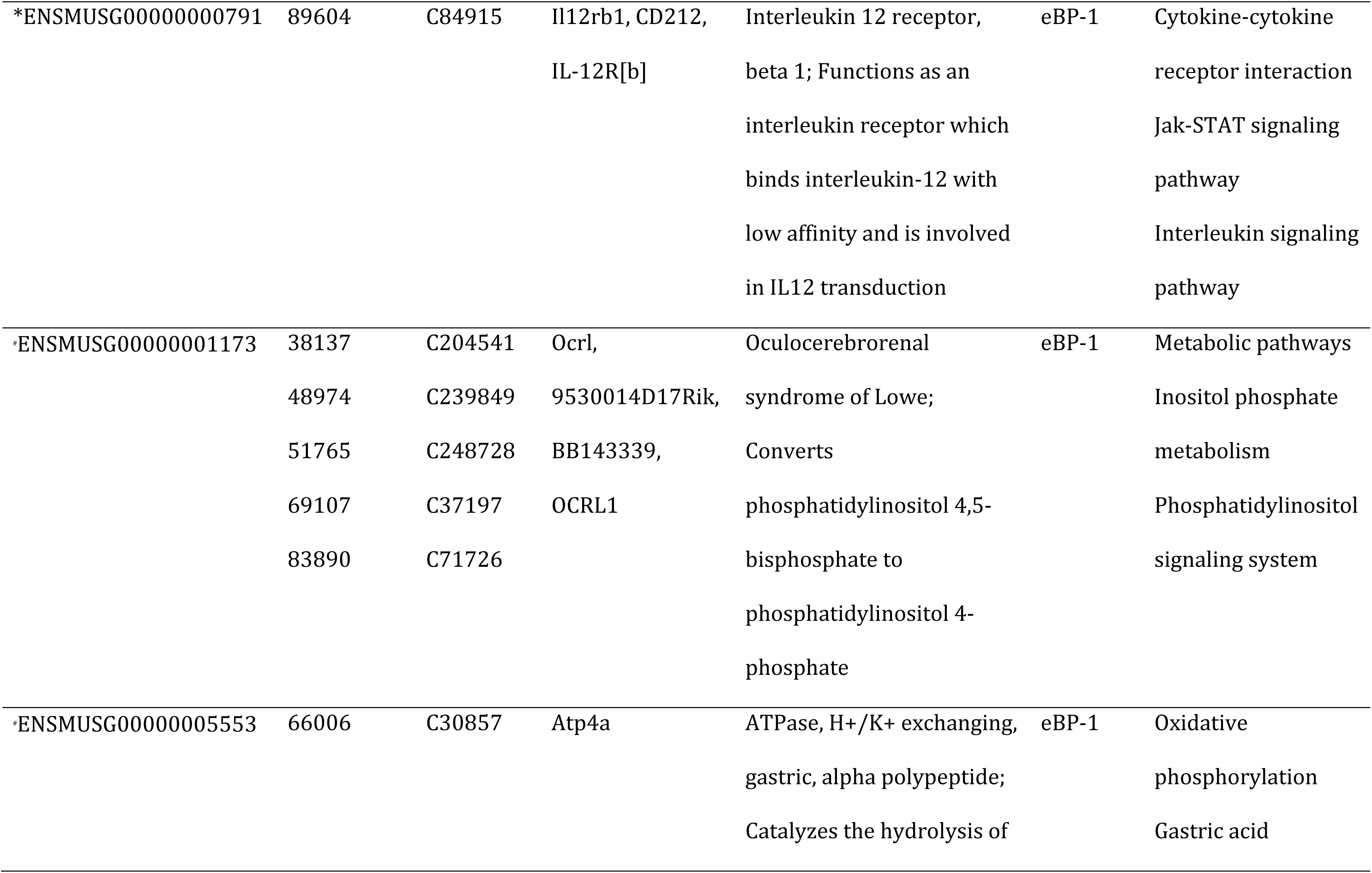
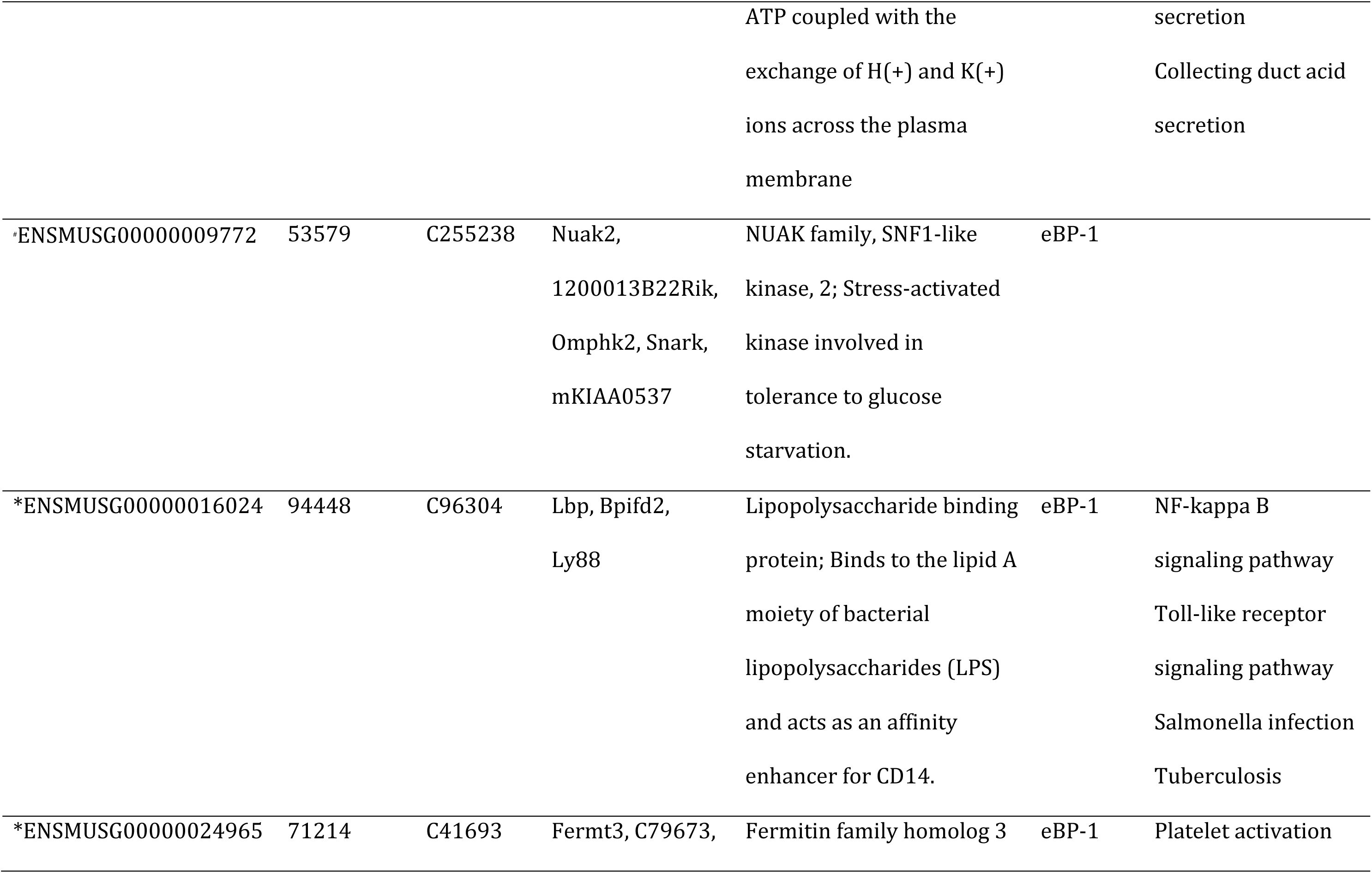
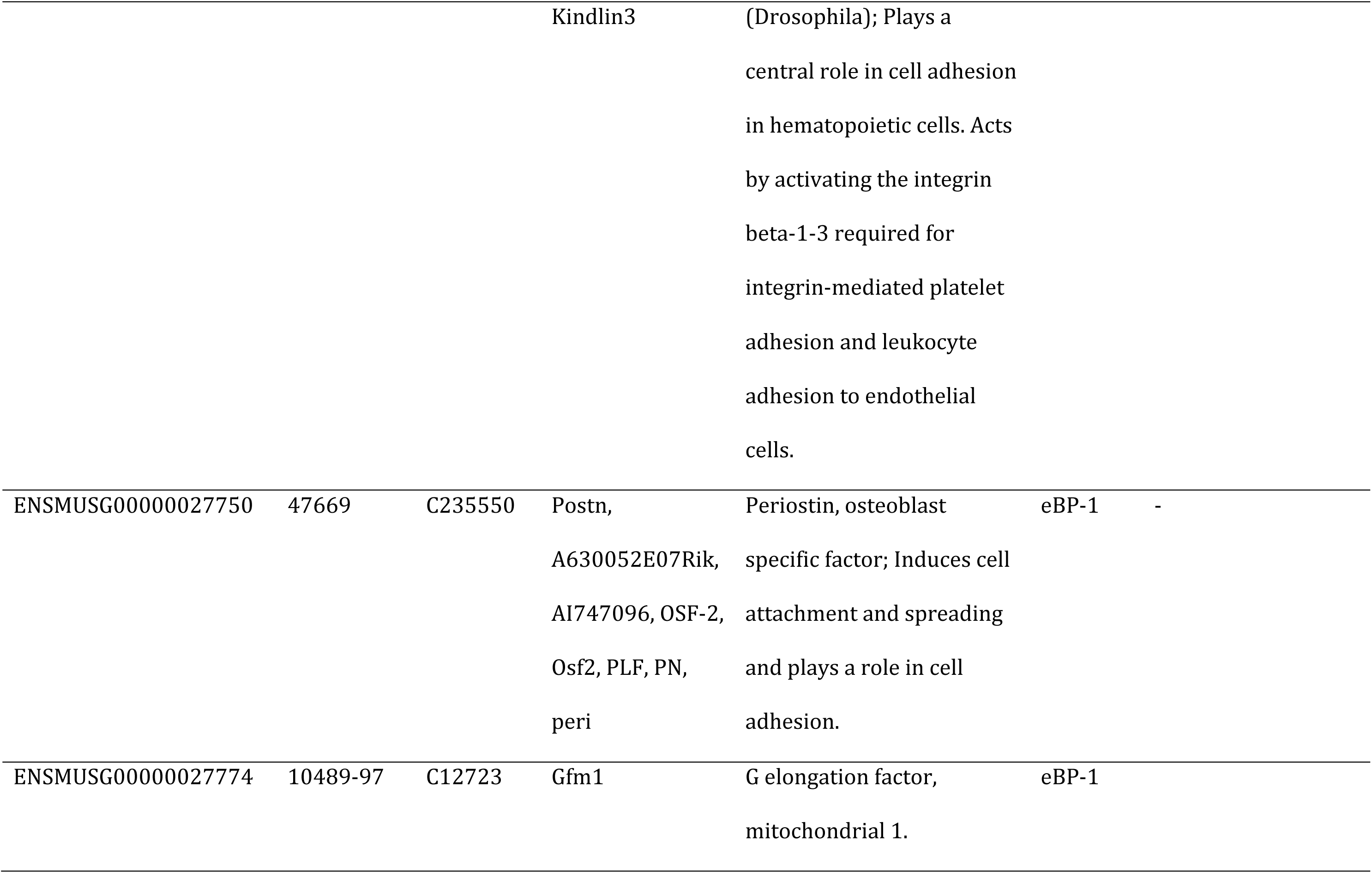
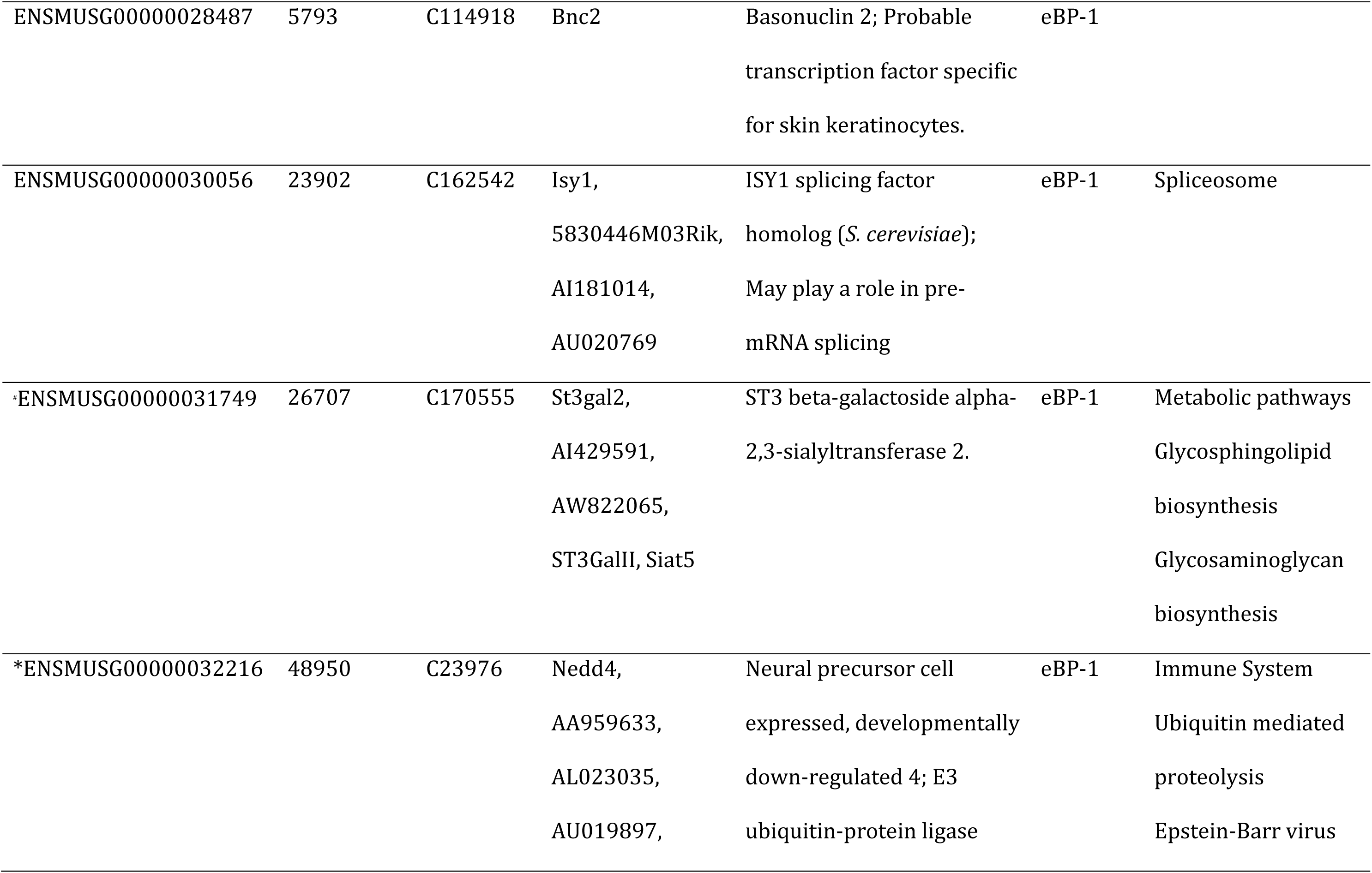
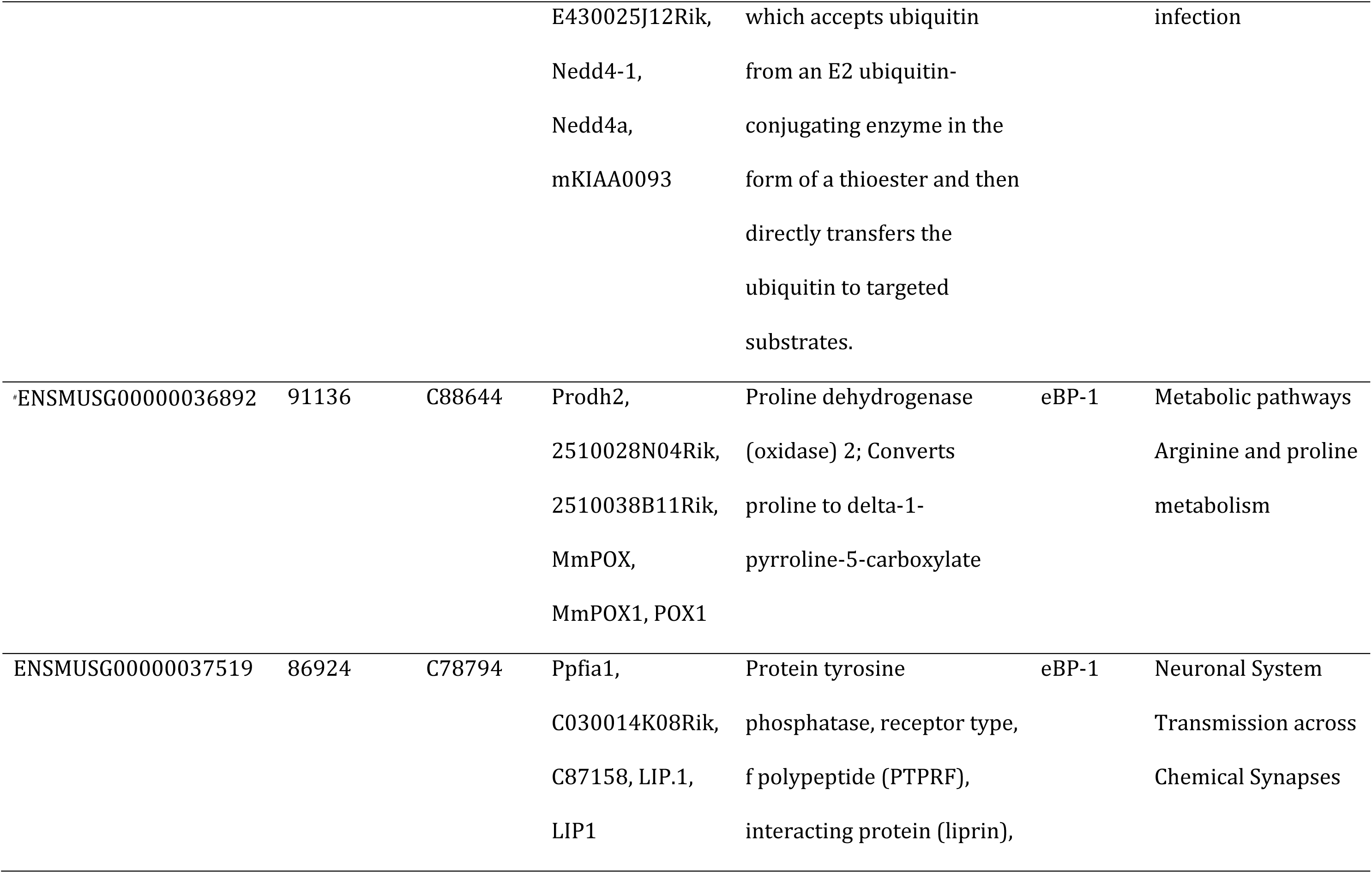
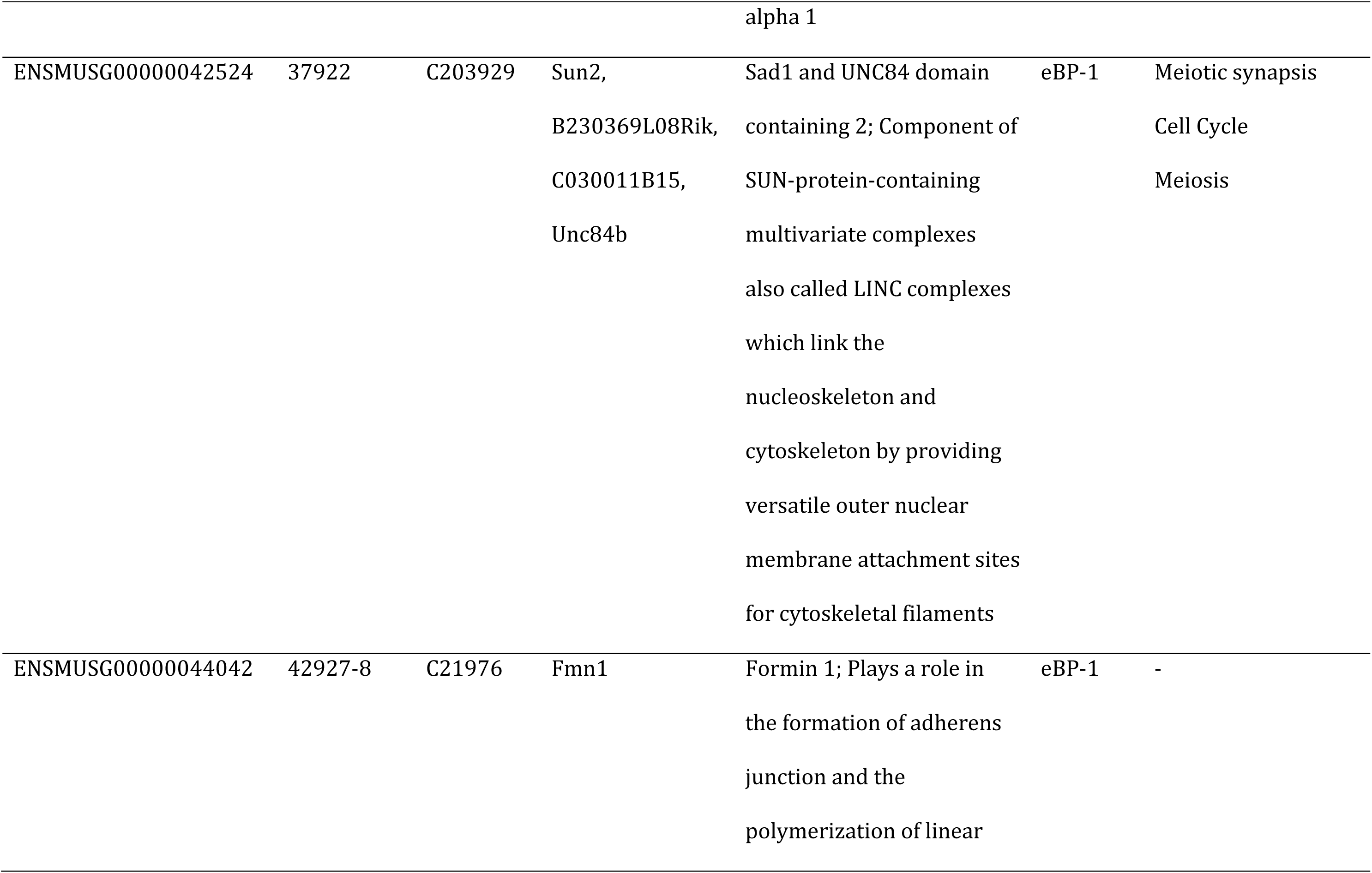
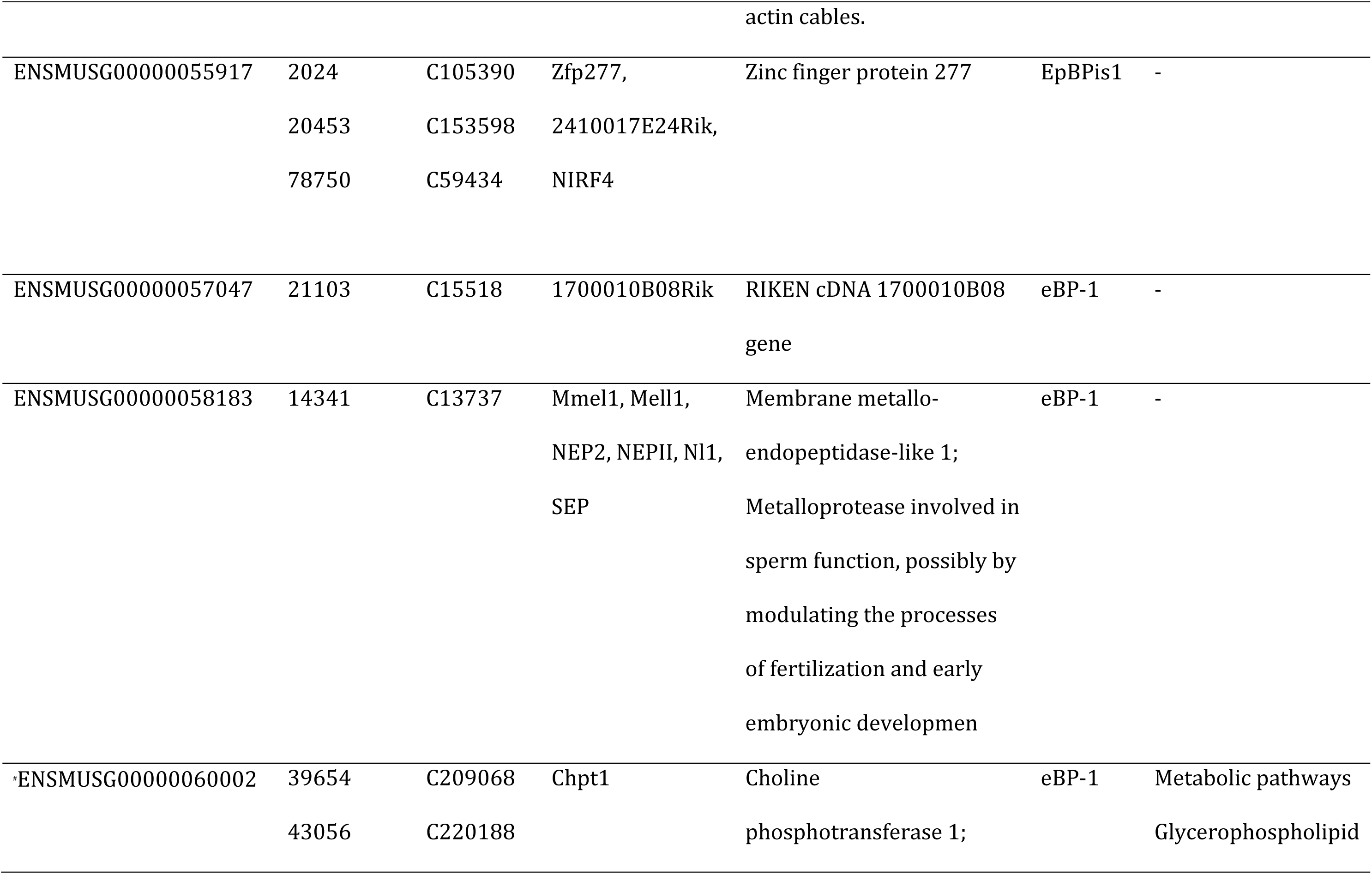
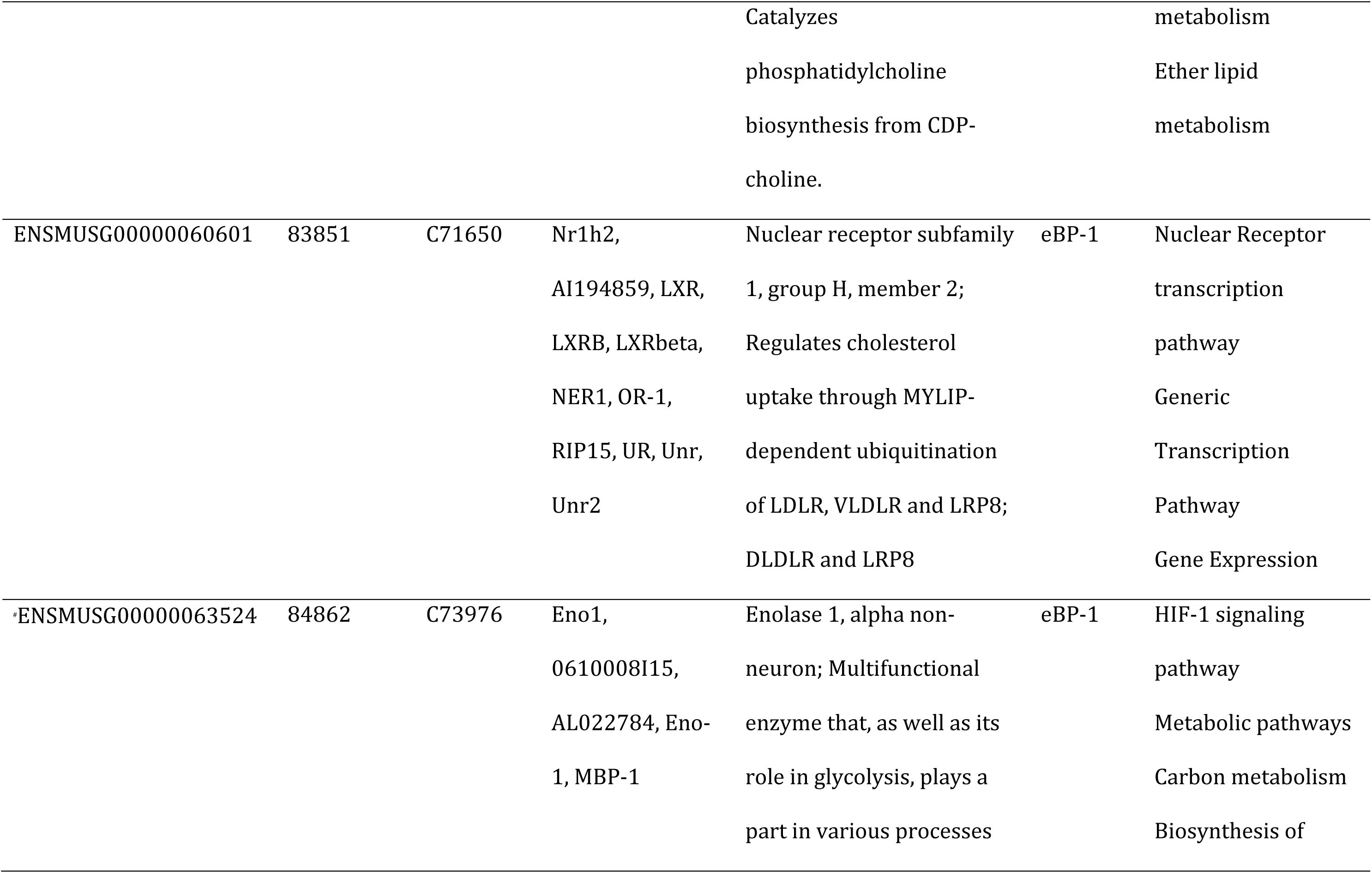
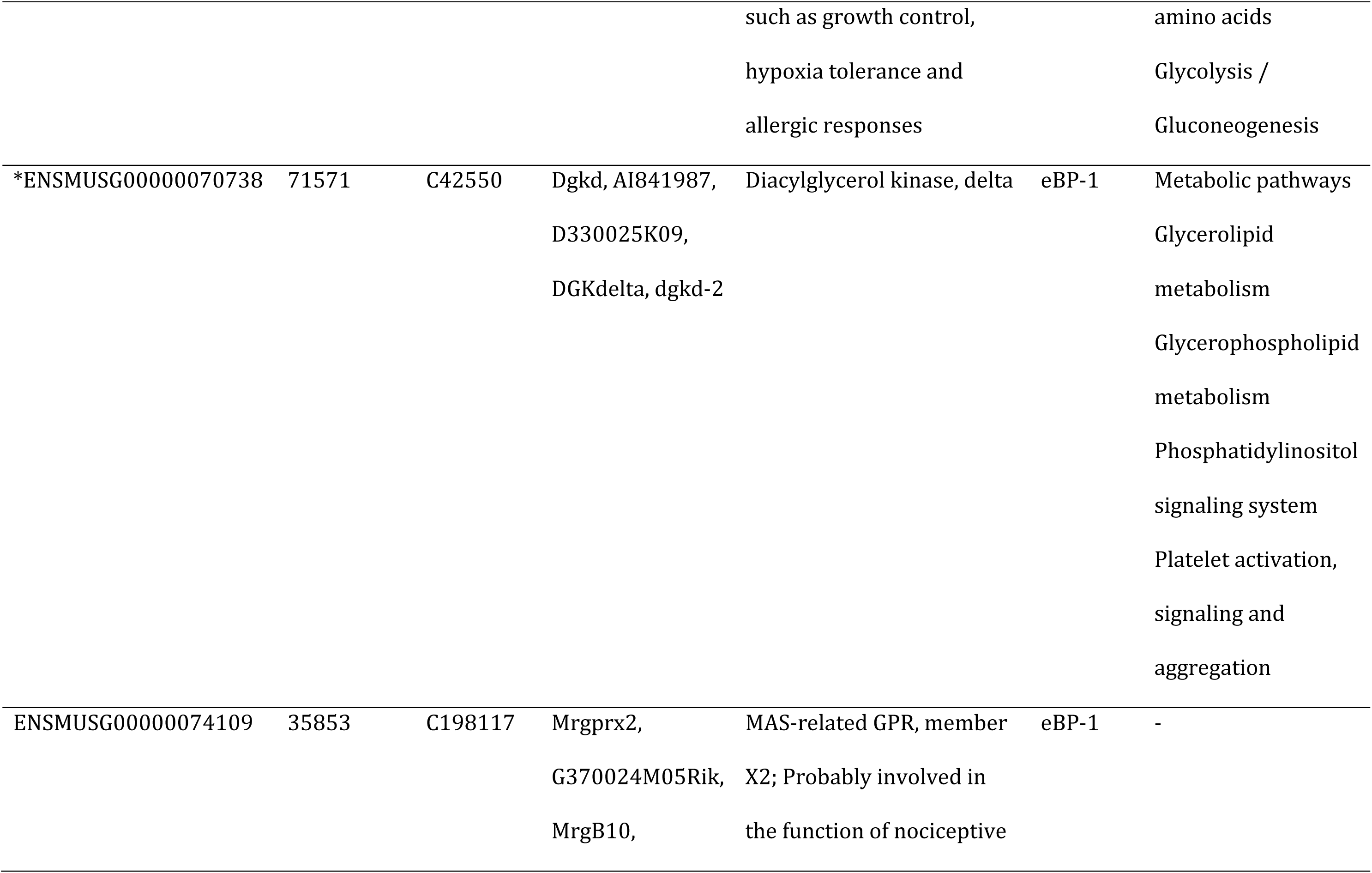
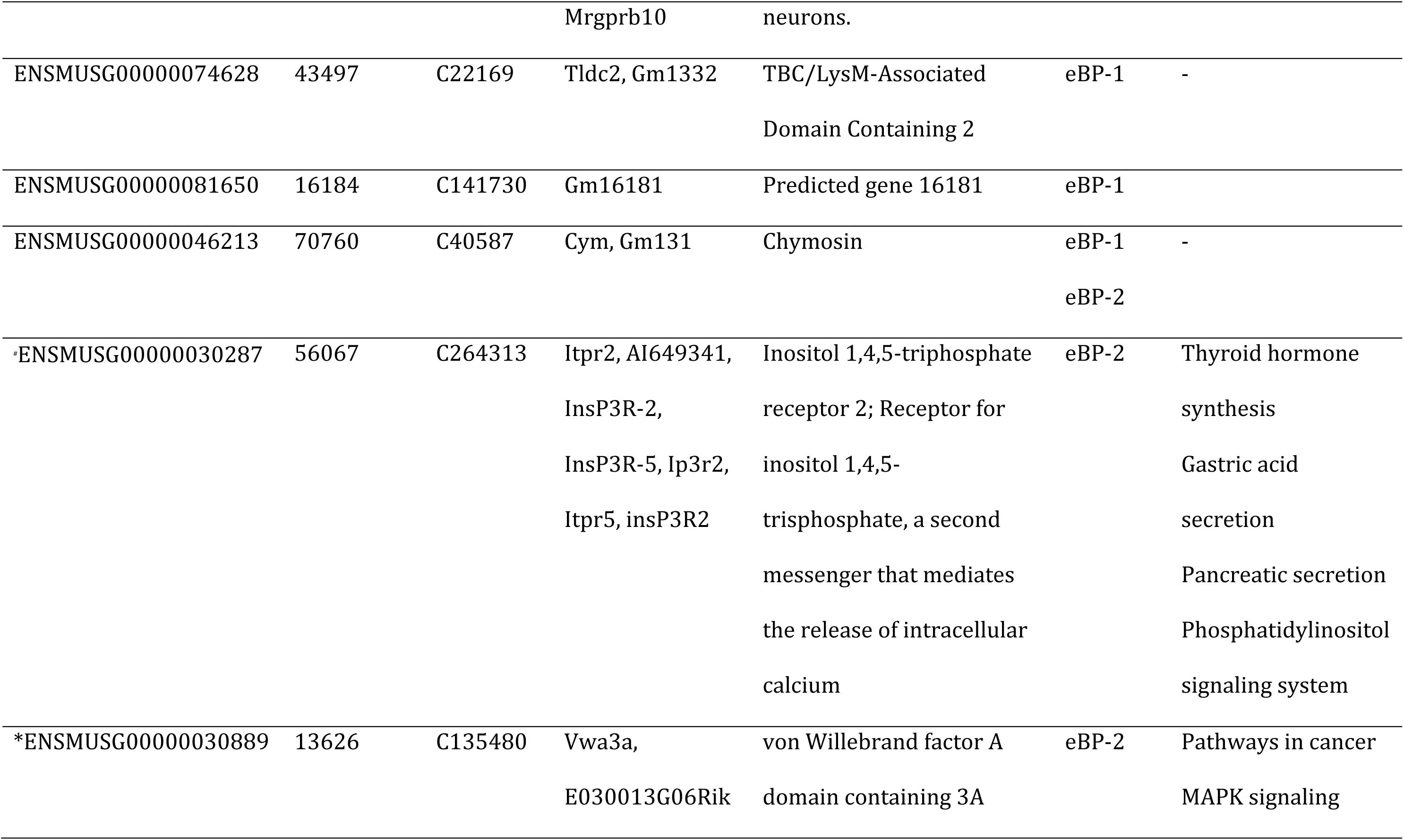
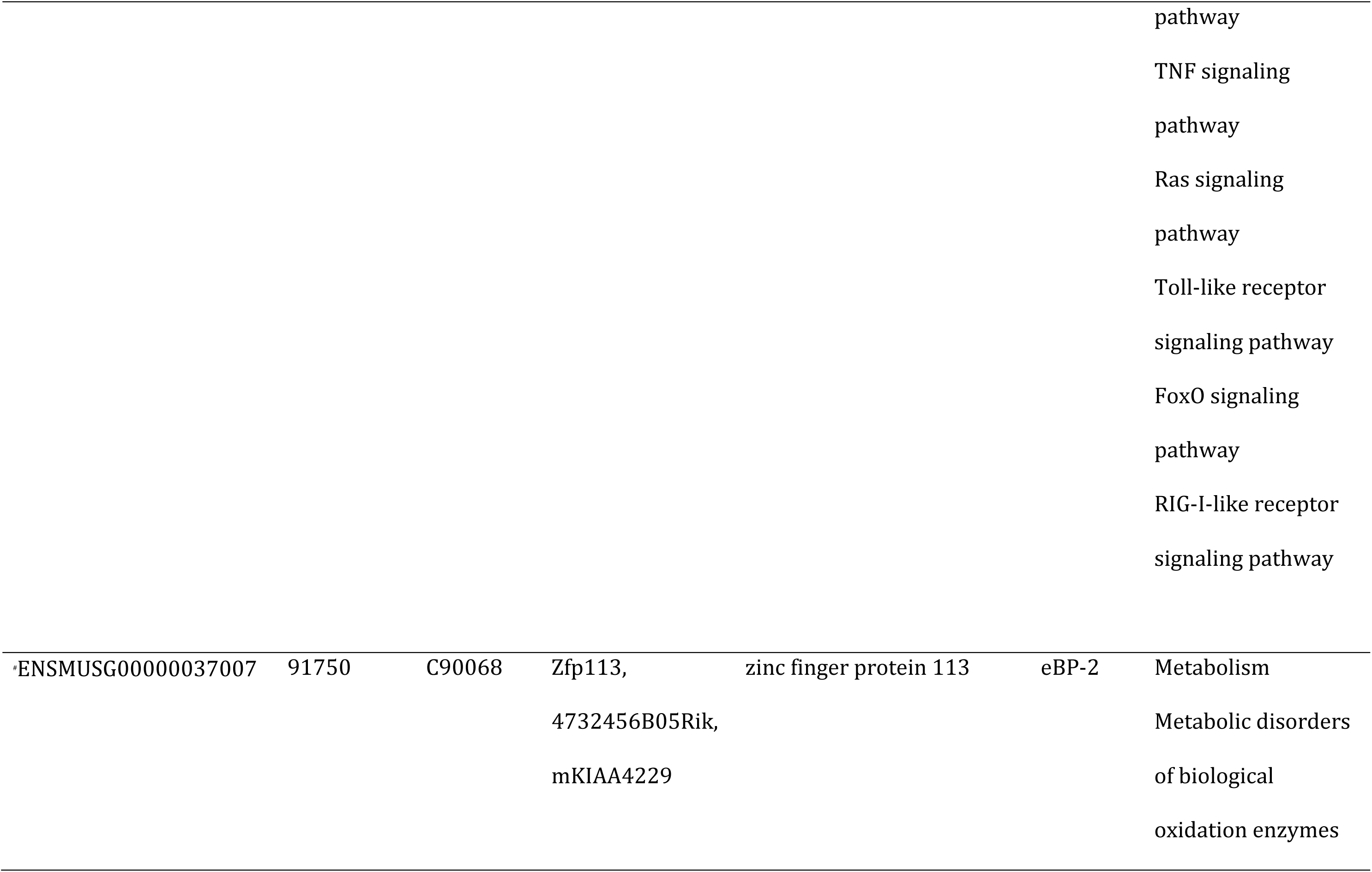
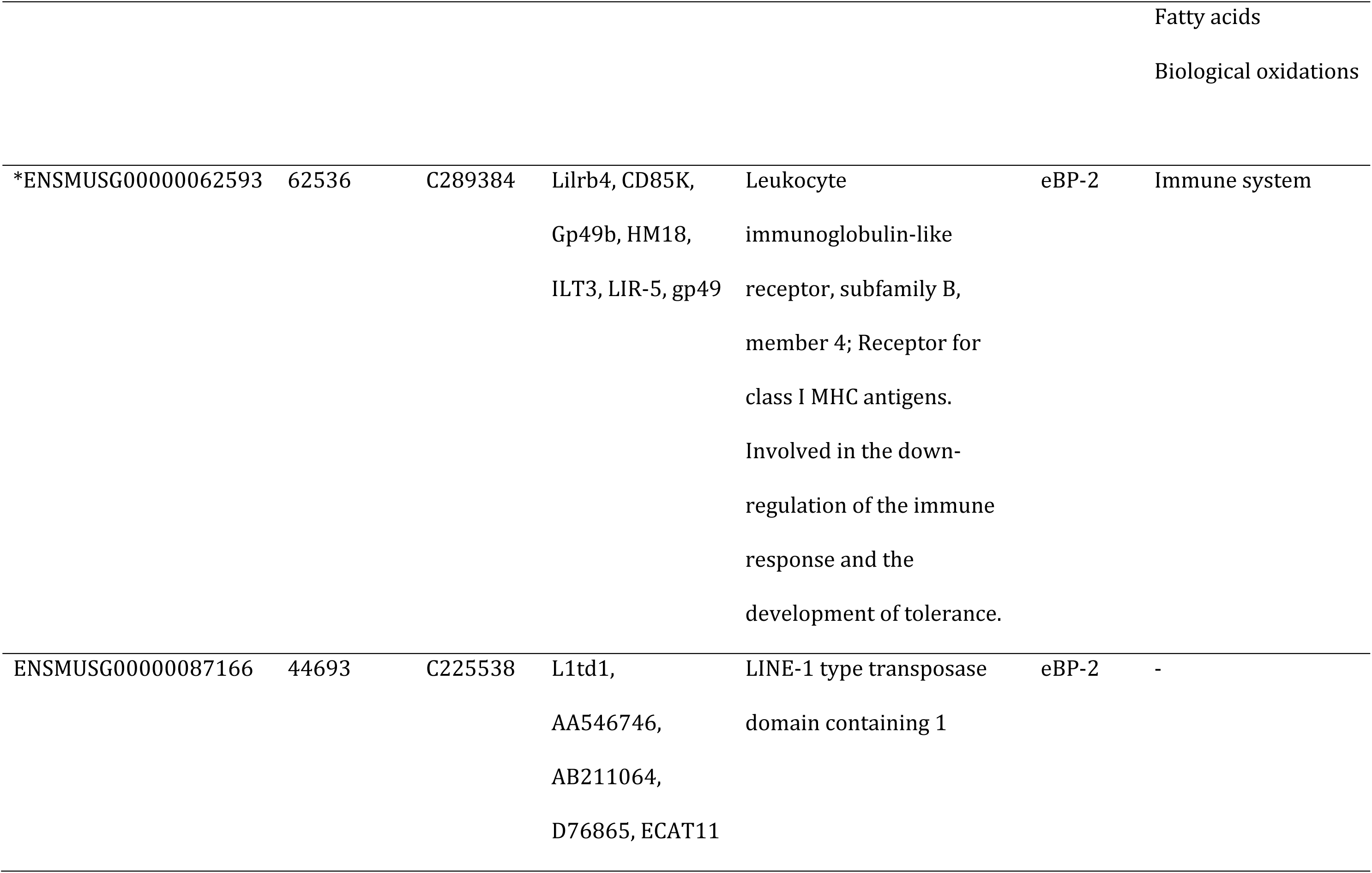
Outlier SNPs identified using at least one of the three methods implemented to detect signatures of selection: SelEstim, BayPass and BayPass with environmental variables (respectively when associations were found with PC1 or PC2). Only SNPS that belonged to contigs that aligned to the mouse genome and corresponded to genes coding for proteins are included here. Gene name and description were obtained in Pathway are indicated following KEGG or Reactom results. Genes related to immunity are indicated by *. Those related to metabolism processes are indicated by^#^.

The protein-protein interaction network analysis emphasized the importance of immunity pathways within this set of annotated outliers. It described eight edges corresponding to associations between five immune related outliers detected in this study (*Ptprc*, *Tlr7*, *Fermt3*, *Dgkd* and *Lilrb4*). The significance of this network (enrichment *p-value* = 0.001) indicated that the encoded proteins are at least partially biologically connected (Fig. 6).

**Fig. 6.**
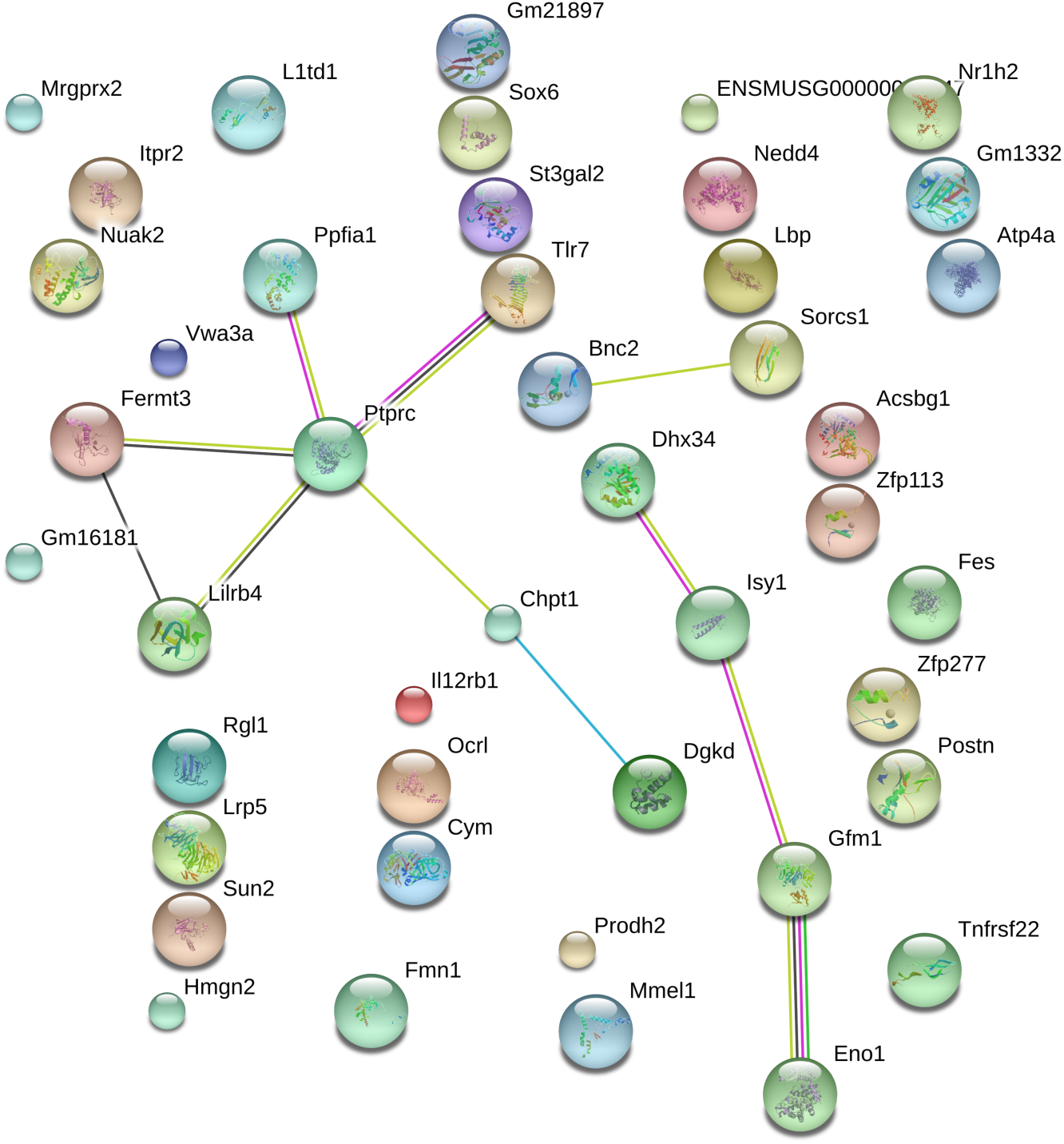
Gene network obtained with STRING including the 52 outliers with annotation based on *M. musculus* genome. Small nodes indicate proteins of unknown 3D structure. Edges represent protein-protein associations based on known interactions (blue: from curated databases; pink: experimentally tested, black= co-expression) or predicted ones (green= gene neighborhood).

Using Kegg pathway, Reactome, PANTHER and BioCyc databases for pathway annotations, we detected 30 pathways with significant enrichment (*p* < 0.05, Table 5). Four of them were linked to TLR pathways, for which we had strong *a priori* reasons to believe that they were involved in adaptive divergence. Seven other significantly enriched pathways were directly related to immunity (‘Antiviral mechanism by IFN-stimulated genes’; ‘Antigen activates B Cell Receptor’; ‘Platelet calcium homeostasis’; ‘citrulline biosynthesis’; ‘ISG15 antiviral mechanism’) or indirectly (‘Elevation of cytosolic Ca2+ levels’; ‘Choline metabolism in cancer’). Another interesting pathway with regard to bank vole / PUUV interactions was ‘Surfactant metabolism’, since some of the proteins involved in this pathway can interact with pulmonary viral infections. All outliers involved in these enriched pathways were detected using the STD model in BayPass, except *Tlr7* that was also detected using SelEstim. Finally, other important classes of enriched pathway categories were related to metabolism, in particular fatty acid metabolism, and neurotransmission.

Among the gene-associated GO terms, 210 had *p*-values below 0.05. Of these GO terms, 176 belonged to the GO category Biological process, 21 to Molecular Function and 13 to Cellular Component. REVIGO formed nine clusters for the category Biological Process (Fig. 7). The most represented cluster was built by the term ‘positive regulation of chemokine production’, followed by ‘substrate adhesion-dependent cell spreading’, ‘myeloid leukocyte activation’ and ‘secretion of lysosomal enzymes’. For the GO category ‘Molecular Function’ nine GO term clusters were formed among which the most common terms comprised ‘beta-galacoside (CMP) alpha-2,3-sialyltransferase activity’, ‘Wnt-activated receptor activity’, ‘phosphoserine binding’ and ‘lipoteichoic acid binding’. Finally, the GO terms ‘receptor complex’ and ‘sarcoplasmic reticulum membrane’ summarised the clusters of the Cellular Component category.

**Fig. 7.**
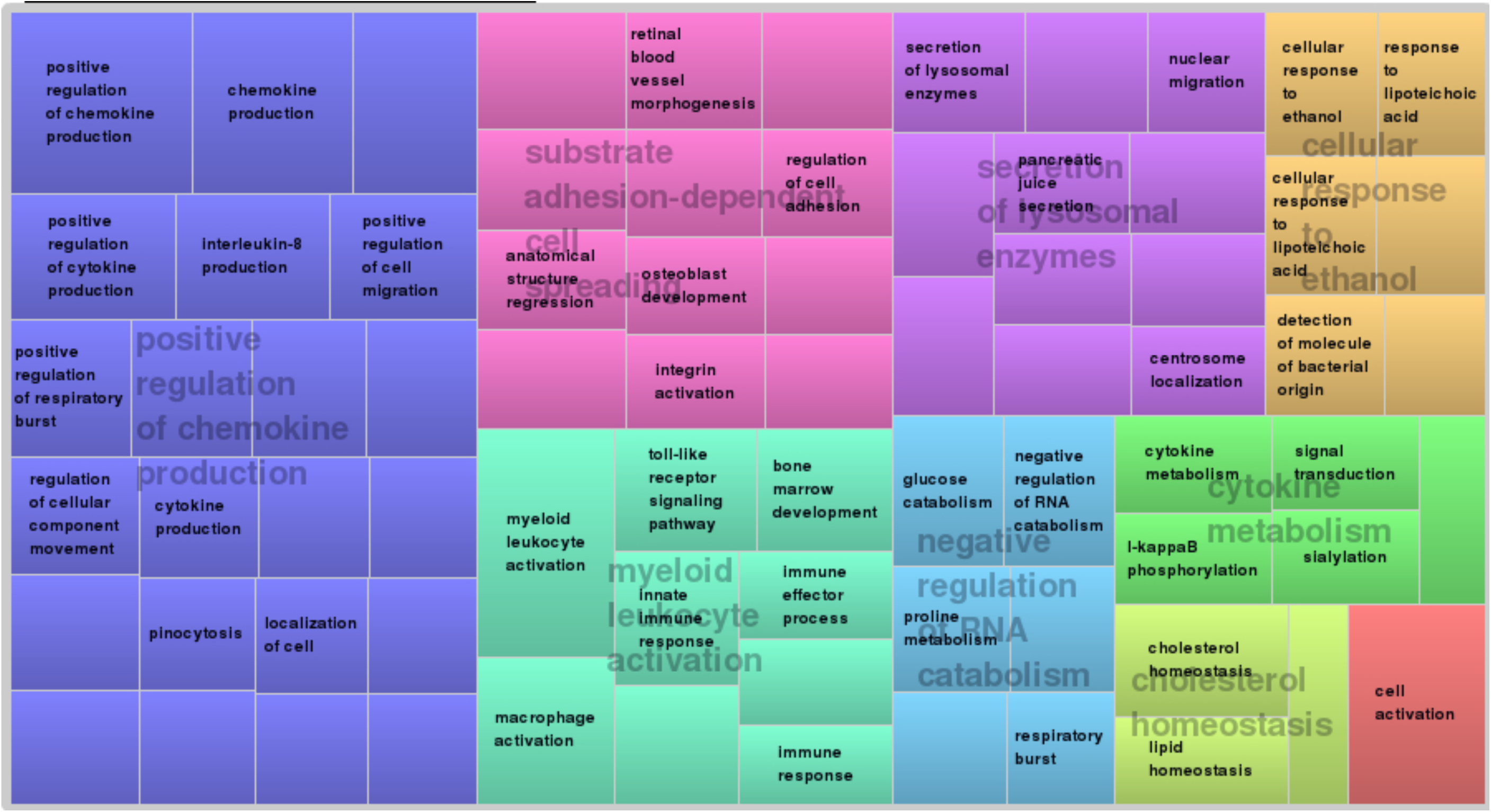
TreeMap view of REVIGO Biological Process analyses. Each rectangle represents a single cluster, that are grouped into ‘superclusters’ of related terms, represented with different colors. The size of the rectangles reflects the frequency of the GO term in the set of outliers included in this analysis.

## Discussion

### A new toolbox for Myodes glareolus studies

Recent years have seen the advent of high throughput sequencing and genotyping technologies and their application to non-model organisms. It has opened new perspectives to perform genomic studies and identify genes and networks involved in a diverse array of ecological and evolutionary processes including speciation, conservation, invasion or biological adaptation (Andrews, Good, Miller, Luikart, & Hohenlohe, 2016; Ekblom & Galindo, 2011; Narum, Buerkle, Davey, Miller, & Hohenlohe, 2013). In the context of zoonoses, these technologies have mostly been used to study newly emerging pathogens (Yang, Yang, Zhou, & Zhao, 2008). The analysis of gene interactions that govern host or reservoir responses to pathogens still remains mostly restricted to laboratory models and major diseases (e.g. malaria vectors, White et al., 2011). In this study, we used paired-end RAD sequencing to examine the genomic patterns of differentiation among six natural populations of *Myodes glareolus,* a rodent reservoir of *Puumala* virus, the agent of a mild hemorrhagic fever with renal syndrom in humans. This study complements previous ones that focused on candidate genes selected from the literature to identify those involved in bank vole susceptibility to PUUV (Charbonnel et al., 2014). It also provides genomic resources of tens of thousands RAD-seq markers that will further be available to study genetic diversity, population structure and adaptation in bank voles. They may be very useful to address different issues related to this rodent in a wide array of disciplines such as medical science (Hampton, 2014; Razzauti-Feliu et al., 2015), microbiology (Kohl, Sadowska, Rudolf, Dearing, & Koteja, 2016), or ecology (Mokkonen et al., 2011).

### High levels of genetic differentiation between northern and southern bank voles

These genomic data provide population structure patterns that are in agreement with the previous phylogeographic studies conducted on bank voles based on mitochondrial sequences and mitogenomes. The stronger level of differentiation was observed between the northernmost population (Hörnefors) and the southern ones (pairwise *F*_ST_ comprised between 0.248 and 0.361), in line with a strong pattern of isolation by distance. This result is also congruent with the macrogeographic pattern classically observed in Fennoscandia, where two main differentiated mitochondrial lineages are present respectively in Southern and North-Eastern Sweden (Jaarola, Tegelstrom, & Fredga, 1999; Tegelstrom & Jaarola, 1998). The recolonization of Fennoscandia, from separate glacial refugia, after the end of the last glaciation period, shaped this phylogeographic pattern, with a southern immigration route that became accessible around 14,1000 BP and a north-eastern one that opened up 10,000 BP (Jaarola et al., 1999). It created a ca. 50 km-wide secondary contact zone between these two mitochondrial lineages, which runs from West to East through Central Sweden, between Hörnefors and Härnösand localities. The genome-wide differentiation observed in this study reflects this phylogeographic history. This contact zone is also detected for other mammalian species including the common shrew, the brown bear and the field voles (Taberlet & Bouvet, 1994). Such suture zone cannot be explained by past or present natural barriers to dispersal (Jaarola et al., 1999), but rather by a secondary contact between divergent recolonizing lineages. Secondary contact might result from similar phylogeographic histories for different mammalian species, or analogous selective pressures acting on these species. Interestingly, Puumala viruses circulate on both sides of this contact zone, with distinct genetic variants in the northern and in the southern bank vole populations, which suggests that a contact zone may also exist between genetically differentiated PUUV lineages (Hörling et al., 1996).

### Biological limitations

Because of the phylogeographical history of *M. glareolus* in Sweden, the transect between the PUUV endemic (Northern Sweden) and non-endemic (Southern Sweden) areas included a contact zone between two genetically differentiated bank vole lineages. Our sampling was therefore marked by a strong genetic structure, that coincided spatially with gradients in ecological variables as well as PUUV distribution.

A first consequence is the possibility that *F*_ST_ outliers do not result from local adaptation but from genetic incompatibilities between different backgrounds. Such endogeneous genetic barriers may result in tension zones, whose locations are initially stochastic but tend to overlap with exogenous ecological barriers (coupling effect, see Bierne, Welch, Loire, Bonhomme, & David, 2011). It is nearly impossible to disentangle *F*_ST_ outliers from genetic incompatibilities in our case, but replicating this work along other PUUV endemic – non-endemic transects could help identifying loci commonly evolving in response to PUUV in bank vole populations.

A second consequence of this sampling strategy concerns the lack of statistical power to detect local adaptation while using *F*_ST_ outlier statistical methods. These methods are often based on theoretical assumptions (e.g., populations at equilibrium, island model of migration) that are rarely completely met in the natural populations sampled. Moreover, spatial autocorrelations in allele frequencies due to isolation by distance can result in false correlations between such frequencies and environmental variables (Lotterhos & Whitlock, 2015; Meirmans, 2012). This potential effect may result in a larger number of false positive detected with SelEstim (which assumes an island model of population structure) as compared to BayPass, and a low number of common outliers between SelEstim and BayPass. This justified the use of genetic-environment associations statistical methods controlling for population structure (here, BayPass). However, this method may also have low statistical power with our sampling design due to the strong correlation between the axis of population genetic differentiation (North-South isolation by distance pattern) and the axis of environmental variations (including variations in PUUV prevalence). Unfortunately, it was not possible to study pairs of nearby populations, as recommended by Lotterhos et al. (2015), because the limit between geographic areas where PUUV pressure is high vs. low is barely known. We therefore had to consider a large sampling scale for bank vole populations, which led to highly genetically structured samples. Finally, it is likely that little information about divergence and selection came from the populations sampled in the middle of the transect, where PUUV pressure as well as environmental and climatic features were intermediate. Future studies should now focus on samples covering the geographic area where clines of allele frequencies exhibit slope disruption.

Because of these biological limitations, genome scan results have to be considered cautiously. The combination of three statistical analyses based on population differentiation and ecological associations to detect outlier loci aimed to reduce the rate of false positives and to enhance our chances of detecting genuine signatures of selection (François, Martins, Caye, & Schoville, 2016). The enrichment tests and gene interaction network analyses may also limit interpretations of false positive outliers.

### Genes involved in selection

We found evidence for a small set of outlier loci (547 SNPs belonging to 468 contigs among 70,699 examined), showing high levels of genetic differentiation consistent with divergent selection acting along our North-South transect in Sweden. Because the whole genome of *Myodes glareolus* is not yet sequenced, assembled and annotated, we met difficulties in annotating RAD contigs and consequently, outlier ones. Only 41 % of the outlier contigs blasted to genes in the mouse genome. Among them, only 21 % belonged to protein coding regions and could be annotated. We therefore could only work on a small part of the information gathered. Moreover we have to remind that this genome scan approach enabled to screen about 35 Mbp, *i.e.* about 1 % of the genome bank vole. Altogether, these facts may explain why this population genomic approach could hardly reveal previously identified candidate genes with regard to bank vole susceptibility to PUUV (Charbonnel et al., 2014). Despite these limits, we have succeeded in identifying other genes and pathways probably evolving under differential selection among bank vole populations from PUUV endemic and non-endemic areas. One third of the enriched pathways representing SNPs showing these signatures of directional selection concerned immune processes, with the ‘positive regulation of chemokine production’ being among the most represented biological process. Infectious pathogens are among the strongest selective forces acting in natural populations, and as such, they contribute to shape patterns of population divergence and local adaptations in the wild through balancing and positive selection acting in host genomes (e.g. Fumagalli et al., 2011; Karlsson et al., 2014; Vatsiou, Bazin, & Gaggiotti, 2016). Many of the immune related genes showing footprint of positive selection here were detected using ecological associations with a synthethic variable describing North / South variations in climatic, forest features and nephropatia epidemica human cases. Although these associations do not reflect causality, they might be biologically meaningful with regard to environmental factors and *M. glareolus* microbiome, including PUUV, that influence bank vole responses to parasitism and genetic polymorphism. Further analyses and experiments are required to confirm and better interpret this result.

More specifically, the analysis of gene interaction network provided support in favor of this potential local adaptation to environment for molecules involved in platelet activation (*Fermt3*, *Dgkd*) and TLR pathway (*Tlr7*). The pattern recognition receptors (PRR) for hantaviruses include Toll-Like receptors, among which TLR7. Its stimulation activates downstream signalling immune cascades with production of pro-inflammatory cytokines and IFNs, which are crucial for inducing a variety of innate antiviral effector mechanisms (Kawai & Akira, 2005). Differential genetic expression of this receptor or different levels of TLR7-Hantavirus recognition due to TLR7 genetic polymorphism could affect this activation of immune cascade, with consequences in terms of virus replication (e.g. for Tlr7 gene expression between sexes, Klein et al., 2004). Moreover, in humans, hantavirus-associated syndromes include increased vascular permeability and platelet dysfunction, with dramatic decreases in platelet counts at the beginning of vascular leakage (Yanagihara & Silverman, 1990). The presence of functional platelet-TLR7 and their activation during viral infections could result in this decrease of viral platelet count, which is likely due to platelet aggregate formation with leukocytes, followed by internalization in the neutrophil population (Koupenova et al., 2014). In addition, hantaviruses bind to α_IIb_β_3_ integrins expressed on platelets and endothelial cells, contributing to viral dissemination, platelet activation, and induction of endothelial cell functions. Gavrilovskaya et al. (2010) suggested that hantavirus associated pathogenesis could be due to the recruitment of quiescent platelets to the surface of infected endothelial cells, thereby forming a platelet covering on the surface of the endothelium. This could dramatically reduce the number of circulating platelets and alter platelet and endothelial cell interactions, which dynamically regulate vascular integrity. In reservoirs, hantaviruses persist without exhibiting any sign of immune pathology and they evade immune responses to establish persistence (Easterbrook & Klein, 2008). Deer mice infected with SNV or ANDV did not show any variations of platelet counts compared to uninfected one {Schountz, 2012 #1409}. Our results might therefore suggest that genetic polymorphisms associated with *Tlr7* or platelet activation genes may account for differences in the immune responses to PUUV infections, and the possibility for this virus to persist in *M. glareolus*. These polymorphisms could contribute to shape the contrasted PUUV epidemiological situations observed in natural populations of bank voles from Northern and Southern Sweden.

In addition, several genes encoding molecules with functions in metabolic processes related to glycolysis / glucogenesis, lipid metabolism and citric acid cycles, showed signatures of selection that were mainly detected from ecological associations with the synthethic variable describing North / South variations in Sweden. Previous studies have demonstrated that energy metabolism shows important inter- and intraspecific variations in endotherms driven by physiological, ecological and evolutionary factors (Boratynski et al., 2011). It is likely that the rate of metabolism is correlated with life history traits and fitness components, although different correlations can be observed due to fluctuating selection between seasons (Nilsson & Nilsson, 2016). We can therefore speculate that polymorphism at these genes (*Acsbg1, Atp4a, Chpt1, Eno1, Hmgn2, Itpr2, Nuak2, Ocrl, Prodh2, St3gal2, Zfp113*, see Table 5) might contribute to bank vole local adaptation to climatic and ecological conditions along a North / South transect accross Sweden. Previous studies based on globin genes in bank voles from Britain found similar patterns of genetic divergence between northern and southern populations, mediated by natural selection through the evolution of bank vole erythrocyte resistance to oxidative stress (Kotlik et al., 2006).

### Methodological considerations

RAD-seq has become an increasingly common genome scan approach this last decade, although several difficulties regarding the identification of genes of functional significance with regard to population divergence and local adaptations have been pointed out (Lowry et al., 2017). Some of these potential limitations are discussed below.

First, we chose to develop a RAD-seq approach based on the pooling of samples without individual barcoding, so that we could estimate population allele frequencies at an affordable cost. Nevertheless, there are several limitations to pooling strategies (Gautier et al., 2013; Narum et al., 2013). Uneven sequencing of individuals in pools may lead to biases in allele frequency estimates. As recommended in Andrews et al. (2016) or Guo et al. (2016), we minimized this bias by including large sample size per pool and replicating libraries for each population. Another possible shortcoming of pooling is that cryptic population structure cannot be identified and taken into account in the case where multiple independent groups of individuals are mixed within a single pool. This possibility was minimized by sampling bank voles in a limited geographic area for each population, although we cannot discard the possibility of intra-population structuring, which is frequent in rodents.

Second, we are aware that the RAD-seq approach enabled to sample only a small proportion of *M. glareolus* genome so that we might have missed potentially important adaptive SNPs (Lowry et al., 2017). This limitation was reinforced by the lack of genomic resources for bank voles, that prevent us from identifying a large part of outlier SNPs. Nevertheless, we think that RAD-seq was the most appropriate approach for bank vole population genomics considering the large genome size of this species, the absence of genome reference and the number of individuals and populations we had to screen (McKinney, Larson, Seeb, & Seeb, 2016).

### Conclusions

Using a pool RAD-seq approch and a combination of statistical methods to detect loci with high genetic differentiation and associations with environmental variables of interests, we have identified a list of putative loci that are worthy for further experimental and functional studies to better understand *Myodes glareolus* / PUUV interactions, and potentially NE epidemiology. These results are of main importance because our previous knowledge of the factors driving immuno-heterogeneity among reservoirs of hantaviruses was inferred from the medical literature or from results based on laboratory models. In the future, they could enable to better assess the risk of NE emergence by including reservoir immunogenetics in ecological niche modelling. They could also have important implications for medical purposes, by revealing potential immune and metabolic pathways driving hantavirus pathogenesis in humans and non-reservoirs.

## Acknowledgments

The authors wish to thank Hélène Holota from TAGC facility for the Covaris sonication of the RAD libraries. Data used in this work were partly produced through the genotyping and sequencing facilities of ISEM (Institut des Sciences de l’Evolution-Montpellier) and Labex Centre Méditerranéen Environnement Biodiversité. This article is registered with the EDENext Steering Committee as EDENext406 (http://www.edenext.eu/).

## Data accessibility

DNA sequences of candidate genes are accessible in Genbank (see accession numbers in Table 3).

Illumina RAD-tag sequences will be accessible in dryad digital depository.

Our final SNP data set is available as a supplementary file.

## Author contribution

A.R., M.Gal. and N.C. designed the research. G. O. performed field sampling. A.R., M.Gal. and K.G. performed the laboratory work. N.C., A.R., B.G., R.V. and M.Gau. helped with bioinformatics and statistical analyses, C.Z. and S.VW provided the environmental data. N.C. wrote the first draft of the manuscript, and all authors contributed substantially to revisions.

